# Target-specific design of drug-like PPI inhibitors via hotspot-guided generative deep learning

**DOI:** 10.1101/2024.10.29.620869

**Authors:** Heqi Sun, Jiayi Li, Yufang Zhang, Shenggeng Lin, Junwei Chen, Hong Tan, Ruixuan Wang, Xueying Mao, Jianwei Zhao, Rongpei Li, Yi Xiong, Dong-Qing Wei

**Affiliations:** State Key Laboratory of Microbial Metabolism, Shanghai-Islamabad-Belgrade Joint Innovation Center on Antibacterial Resistances, Joint International Research Laboratory of Metabolic & Developmental Sciences and School of Life Sciences and Biotechnology, Shanghai Jiao Tong University, Shanghai, 200240, China; Qihe Laboratory, Qishui Guang East, Qibin District, Hebi, Henan, 458030, P.R. China; School of Mathematical Sciences and SJTU-Yale Joint Center for Biostatistics and Data Science, Shanghai Jiao Tong University, Shanghai 200240, China; Zhongjing Research and Industrialization Institute of Chinese Medicine, Zhongguancun Scientific Park, Meixi, Nanyang, Henan, 473006, China; Shanghai Innovation Institute, Shanghai, China; Artificial Intelligence Biomedical Center, Zhangjiang Institute for Advanced Study, Shanghai Jiao Tong University, Shanghai, 201203, China; Peng Cheng National Laboratory, Shenzhen, 518055, China

## Abstract

Protein–protein interactions (PPIs) are vital therapeutic targets. However, the large and flat PPI interfaces pose challenges for the development of small-molecule inhibitors. Traditional computer-aided drug design approaches typically rely on pre-existing libraries or expert knowledge, limiting the exploration of novel chemical spaces needed for effective PPI inhibition. To overcome these limitations, we introduce Hot2Mol, a deep learning framework for the de novo design of drug-like, target-specific PPI inhibitors. Hot2Mol generates small molecules by mimicking the pharmacophoric features of hot-spot residues, enabling precise targeting of PPI interfaces without the need for bioactive ligands. The framework integrates three key components: a conditional transformer for pharmacophore-guided, drug-likeness-constrained molecular generation; an E(n)-equivariant graph neural network for accurate alignment with PPI hot-spot pharmacophores; and a variational autoencoder for generating novel and diverse molecular structures. Experimental evaluations demonstrate that Hot2Mol outperforms baseline models across multiple metrics, including docking affinities, drug-likenesses, synthetic accessibility, validity, uniqueness, and novelty. Furthermore, molecular dynamics simulations confirm the good binding stability of the generated molecules. Case studies underscore Hot2Mol’s ability to design high-affinity and selective PPI inhibitors, demonstrating its potential to accelerate rational PPI drug discovery.

## Introduction

Protein–protein interactions (PPIs) are crucial in the regulation of numerous biological functions. Dysrégulation of PPIs can lead to various diseases, including cancer, infectious diseases, and neurodegenerative disorders ^1–3^. Targeting PPIs offers advantages such as high selectivity, reduced off-target effects, and lower chances of resistance compared to targeting single proteins or enzymes ^4,5^. Several PPI inhibitors have already been approved by the FDA ^6^, with small molecules standing out due to their superior cell permeability, oral bioavailability, and structural stability compared to peptides and antibodies. However, targeting PPIs with small molecules is challenging due to the structural characteristics of PPI targets. PPI interfaces are typically large (1,500–3,000 Å^2^) ^7^, which are much bigger than receptor-ligand interactions (300–1,000 Å^2^) ^8^, and are predominantly hydrophobic ^7^. Additionally, they often lack distinct grooves or pockets, making effective binding by small molecules difficult.

Despite the complexity of PPI interfaces, identifying hot-spots ^9^ provides a focus for the rational design of PPI inhibitors. Hot-spot residues contribute significantly to binding energy, allowing small molecules to efficiently bind and disrupt protein interactions. Two main strategies are commonly used in designing small-molecule inhibitors targeting these hot-spots. The first strategy involves molecular docking to identify fragments that bind to hot-spots, which are then optimized into lead compounds ^10^. The second approach designs small molecules to mimic the pharmacophore of hot-spot residues, allowing them to bind effectively to complementary regions on the target protein. A pharmacophore refers to a set of spatially arranged chemical features that are crucial for molecular interactions. This strategy is exemplified by Idasanutlin, which disrupts the MDM2/p^53^ interaction by mimicking the aromatic side chain of Trp23 and the hydrophobic side chains of Leu26 and Phe19, which are critical for binding to MDM2 ^11^. This latter strategy often leads to higher specificity and affinity, as it more closely mimics natural protein-protein interactions.

Traditional computer-aided drug design (CADD) methods have been developed to successfully identify many potent drug candidates, but they are often limited to existing virtual libraries or expert knowledge ^12–14^. In contrast, deep generative models (DGMs) are emerging as powerful tools that explore vast chemical spaces to generate novel compounds and optimize them for desired properties. Traditionally, DGMs for molecular generation have been predominantly ligand-based ^15–18^. These models use neural networks to learn a probability distribution of structural features from known bioactive compounds and subsequently sample novel molecular structures from the learned distributions. However, a major limitation of ligand-based DGMs is their neglect of the role of target protein structure. Recent advancements have led to the development of structure-based DGMs, which integrate explicit protein structural information into the generation process ^19–23^. This integration is crucial, as the efficacy of a drug molecule largely depends on its precise binding to the target. Additionally, many protein targets have few or no known ligands, which makes ligand-based DGMs less generalizable. Structure-based DGMs are expected to overcome these drawbacks and are attracting growing attention.

Despite the potential of structure-based DGMs to generate novel and high-affinity molecules for specific protein pockets, their application in PPI-targeted drug design remains limited. Most existing DGMs are designed for enzymes or transcription factors, which possesses observed ligand-binding cavities or pockets. In contrast, few DGMs are specifically designed for PPIs in lack of deep pockets and these are predominantly ligand-based methods ^18,24,25^. Moreover, while conventional structure-based DGMs excel at ensuring the geometric accuracy and chemical validity of generated molecules, they often do not explicitly optimize molecular properties such as drug-likeness, which are vital for successful drug development. To advance this field, it is essential to develop structure-based DGMs that incorporate the unique characteristics of PPI structures while optimizing molecular properties to design high-affinity and drug-like inhibitors.

In this work, we introduce Hot2Mol, a deep learning model specifically designed for the de novo generation of target-specific, drug-like PPI inhibitors (Fig. 1). Hot2Mol addresses the challenges of targeting large and flat PPI interfaces by generating small molecules that mimic the pharmacophoric features of hot-spot residues, enabling precise inhibition of these interactions. At its core, Hot2Mol integrates a conditional transformer with a variational autoencoder (VAE). The transformer encoder leverages attention mechanisms to capture the intricate relationships between molecular structures, pharmacophoric features, and property constraints. E(n)-equivariant graph neural networks (EGNNs) are employed to encode molecular structures and pharmacophoric features in 3D space, ensuring that spatial information is captured accurately and geometric equivariance is preserved. The VAE processes the transformer’s encoded data, mapping it into a shared latent space from which it samples diverse latent variables. These latent variables guide the autoregressive transformer decoder to reconstruct molecules in SMILES format, ensuring they meet the specified pharmacophoric and drug-like property conditions.

**Fig. 1.**
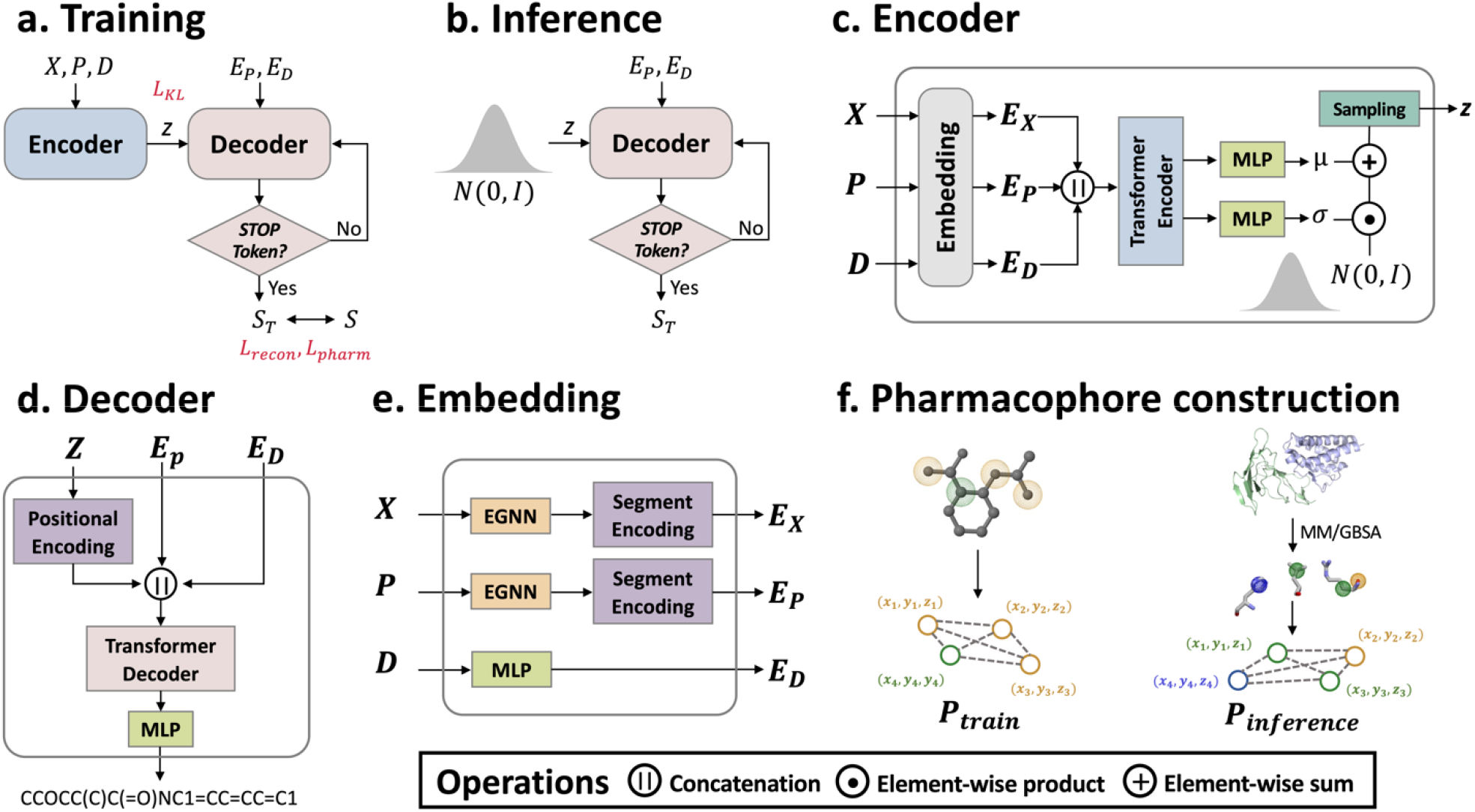
Overview of Hot2Mol. (**a**) **Training phase:** Inputs include molecular graph (*X*), pharmacophore graph (*P*), and molecular property constraint (D). The target molecule is autoregressively reconstructed in SMILES format, minimizing,, and. (**b**) **Inference phase:** A latent vector *z* is sampled from a standard normal distribution and combined with pharmacophore embeddings () and property constraint embeddings () to generate target molecules, (**c**) **Encoder module:** Embeds all inputs into a shared latent space using the latent vector *z*. (**d**) **Decoder module:** Autoregressively generates a molecule in SMILES format from the latent vector *z*, incorporating and. (**e**) **Embedding module:** Molecular graph *X* and pharmacophore graph *P* are embedded using E(n)-equivariant graph neural networks to obtain and. Property encoding *D* is embedded to obtain. (**f**) **Pharmacophore construction:** During training, pharmacophores are randomly selected to build. During inference, pharmacophores are sampled from hot-spot residues of the PPI target protein to build.

Our experiments show that Hot2Mol generate molecules with binding energy distributions superior to both experimentally validated inhibitors and those produced by state-of-the-art DGMs. We further evaluated our model by assessing vital properties of generated molecules, including binding stability, novelty, PPI-targeting drug-likeness, synthetic accessibility, chemical space distributions, and geometric patterns. To illustrate Hot2Mol’s practical applicability in structure-based drug design, we conducted case studies showing its ability to generate high affinity and selective inhibitors for therapeutically relevant targets.

## Results

### Evaluation of common properties for generated molecules

We evaluated Hot2Mol by analyzing several common metrics of the generated molecules, including validity, uniqueness, novelty, the ratio of available molecules, synthetic accessibility (SA), and quantitative estimate of PPI targeting drug-likeness (QEPPI). The definitions of these metrics are provided in Methods section. For comparative analysis, we benchmarked Hot2Mol against three state-of-the-art molecular DGMs: Pocket2Mol ^23^, Lingo3DMol ^26^, and PGMG ^27^ Pocket2Mol is a structure-based DGM that generates molecules conditioned on the 3D geometry of protein pockets. Lingo3DMol extends this approach by combining language models with geometric deep learning to generate 3D molecules based on pocket structures. We selected these two structure-based DGMs as benchmarks to highlight how Hot2Mol offers a more effective solution by targeting hot-spot residues in PPI interfaces. PGMG, which also employs a pharmacophore-guided approach, was included as a relevant model for comparison.

These models were evaluated on a test set of ten randomly selected PPI targets from the DLiP database ^28^, a curated database of PPI modulators. These targets include MENIN/MLL (PDB: 3U85), IL-2/IL-2R (PDB: 1Z92), KRAS/SOS1 (PDB: 6EPM), CBP/H3 (PDB: 5GH9), ZIPA/FTSZ (PDB: 1F47), XIAP/CASPASE-9 (PDB: 1NW9), BCL2/BAX (PDB: 2XA0), MDM2/P53 (PDB: 1YCR), TCF4/Beta-catenin (PDB: 1JDH), and BRD4/H4 (PDB: 3UVW). For each target, 1,000 molecules were generated by using each DGM.

In sum, the 10,000 molecules were generated for the ten targets by using each DGM. The average values of six properties for molecules generated by each DGM are presented in Table 1, with detailed results in Supplementary Tables 1-4. We prioritized the ratio of available molecules as a key metric derived from validity, novelty, and uniqueness scores. Table 1 shows that Hot2Mol outperforms the three baseline models in the ratio of available molecules, demonstrating superior capability in generating valid and unique molecules. As demonstrated in the Ablation Studies section, this performance is largely attributed to the use of EGNNs, which enhance the accurate representation of molecular structures and pharmacophore distributions.

**Table 1.**
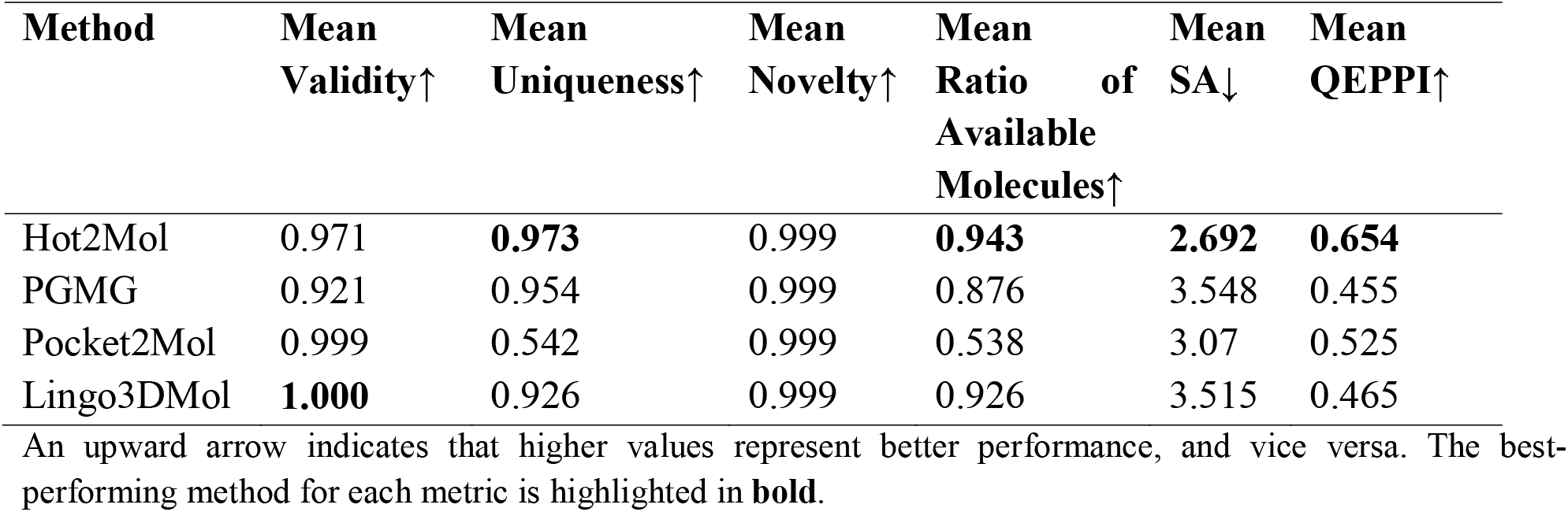
Comparison of generated molecules by four DGMs for ten PPI targets.

Moreover, Hot2Mol achieves a QEPPI value of 0.654, significantly outperforming the three baseline methods (0.455 for PGMG, 0.525 for Pocket2Mol, and 0.465 for Lingo3DMol), indicating its superior efficacy in generating drug-like molecules. This advantage arises from the conditional transformer’s ability to generate molecules under drug-like property constraints, as validated in the Ablation Studies section. Furthermore, the SA scores indicate that Hot2Mol-generated molecules, with an average SA score of 2.692, are easier to synthesize compared to those from baseline models, which have scores ranging from 3.07 to 3.548. Additionally, most Hot2Mol-generated molecules meet pharmacokinetic and toxicity criteria for drug candidates, according to the guidelines of ADMET lab 3.0 ^29^ (Supplementary Fig. S1), demonstrating that Hot2Mol can generate PPI inhibitors with favorable ADMET profiles.

### Analysis of Hot2Mol on chemical space distribution

To assess the chemical space distribution of Hot2Mol-generated molecules, we compared them with known bioactive molecules for the same ten PPI targets. The bioactive molecules were sourced from the DLiP ^28^ databases. A set of 5,000 molecules was randomly selected from both the bioactive and Hot2Mol-generated molecule sets. Given the importance of both substructural and 3D characteristics in molecular design, we employed MACCS ^30^, RDKit ^31^, and USRCAT (Ultrafast Shape Recognition with CREDO Atom Types) ^32^ fingerprints to represent the chemical space of molecules. MACCS fingerprints capture predefined structural keys like functional groups, while RDKit fingerprints encode 2D substructures based on atom and bond types. USRCAT enhances the USR (Ultrafast Shape Recognition) algorithm by incorporating pharmacophoric information to capture the molecular 3D shape.

The chemical space distribution, visualized using Uniform Manifold Approximation and Projection (UMAP) (Fig. 2), shows that Hot2Mol-generated molecules exhibit significant overlap with bioactive molecules in both 2D and 3D chemical spaces, indicating strong structural and pharmacophoric similarity. Similarly, the molecules generated by the three baseline DGMs (Supplementary Fig. S2-S4) also display a comparable level of overlap with bioactive compounds in the same spaces. This result suggests that all models, including Hot2Mol, are effective in generating molecules that occupy relevant regions of the chemical space, ensuring structural alignment with known bioactive compounds.

**Fig. 2.**
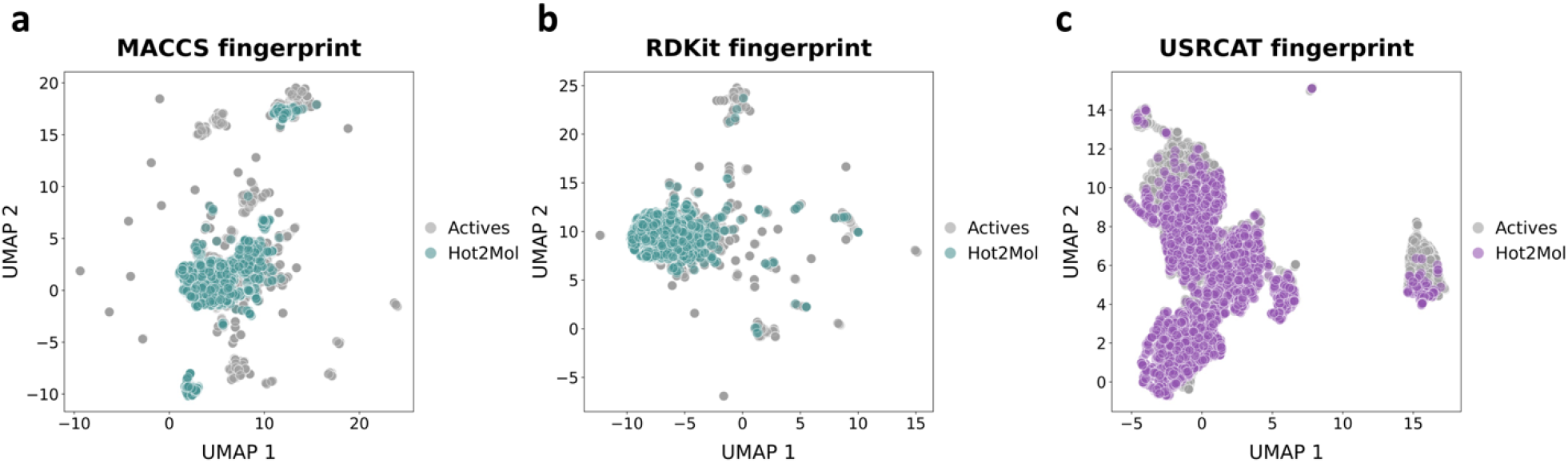
Visualization of chemical space distribution. (**a**) MACCS fingerprints, (**b**) RDKit fingerprints, and (**c**) USRCAT fingerprints visualized using UMAP in two-dimensional space. MACCS and RDKit fingerprints encode sub-structural features, whereas USRCAT fingerprints encode 3D chemical structure.

### Analysis of geometric distribution of generated molecules

We analyzed the ring size distribution for molecules generated by Hot2Mol, those in the training set, and known bioactive molecules (Fig. 3a). The mean ring size for Hot2Mol-generated molecules is 5.77, which closely matches that of the training set (5.73) and bioactive molecules (5.75). Most rings generated by Hot2Mol are 5-membered or 6-membered, which are important for synthetic accessibility, chemical stability, and pharmacokinetic properties. This consistency in ring size suggests that Hot2Mol-generated molecules share key characteristics with bioactive compounds and the training set.

**Fig. 3.**
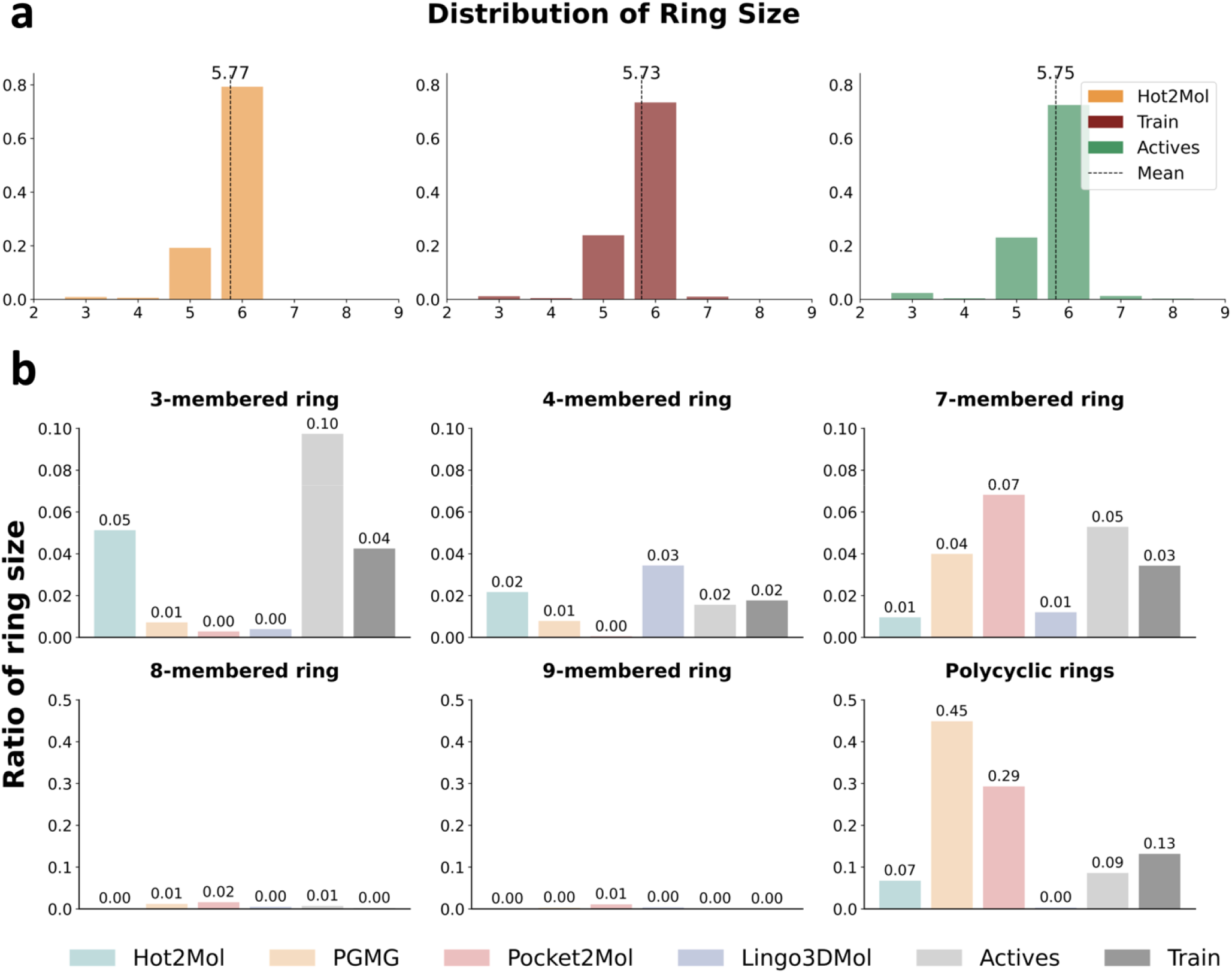
Evaluation of the geometry of generated molecules. (**a**) Distribution of ring sizes in molecules generated by Hot2Mol, molecules from the training set, and bioactive molecules. (**b**) Proportion of molecules containing rings of various sizes, generated by Hot2Mol and three baseline methods, and training and bioactive molecules.

For a more detailed analysis, we examined the proportion of molecules containing rings of various sizes. Drug-like molecules typically favor 5- and 6-membered rings. Conversely, rings with three, four, seven, or more members, including polycyclic rings (three or more fused rings), are less common due to challenges such as low chemical stability, high toxicity, and metabolic instability. Therefore, we particularly checked the occurrence of 3-, 4-, 7-, 8-, and 9-membered rings, as well as polycyclic rings (Fig. 3b). Hot2Mol and Lingo3DMol generated very few unstable macro rings (7-, 8-, and 9-membered), each comprising less than 1% of the molecules. In contrast, Pocket2Mol and PGMG produced 10% and 5% macro rings, respectively. Additionally, Hot2Mol generated fewer polycyclic rings (7%) compared to bioactive molecules (9%) and the training set (13%). In contrast, PGMG and Pocket2Mol had much higher proportions of polycyclic rings at 45% and 29%, respectively.

Hot2Mol generated a higher proportion of 3-membered (5%) and 4-membered rings (2%) compared to PGMG and Pocket2Mol, which produced fewer than 1% of these smaller rings. While these small rings are generally less favorable, they are still found in real-world datasets, with bioactive molecules and the training set containing up to 10% 3-membered rings and 2% 4-membered rings. The presence of smaller rings in Hot2Mol-generated molecules demonstrates the model’s ability to capture diverse ring structures, balancing stability and synthetic feasibility with less common yet pharmacologically interesting designs. Additionally, we evaluated the bond lengths and angles in these molecules, covering nine common covalent bond types (C-C, C-Cl, C-F, C-N, C-O, C-S, C=C, C=N, C=O) (Supplementary Fig. S5) and eight common bond angles (CCC, C=CC, CN=C, CNC, COC, CSC, NCC, O=CO) (Supplementary Fig. S6). The results show that Hot2Mol-generated molecules exhibited distributions consistent with both the training set and bioactive molecules.

### Binding affinity analysis of de novo designed molecules

Beyond geometric structures, we further investigated the interaction strength between ligands and PPI interfaces. Utilizing both the generated molecules and known bioactive compounds, we employed AutoDock Vina ^33^ to approximate binding affinities between ligands and proteins. The protein PDB structures used for docking are listed in Methods section. To mitigate the inherent bias towards higher binding affinities associated with larger molecules—stemming from their potential to engage in more interactions with protein—we restricted our analysis to ligands with molecular weights below 600 Da.

Fig. 4a illustrates the docking score distributions for both generated and bioactive molecules. Hot2Mol-generated molecules exhibited docking scores comparable to, or even better than those of bioactive molecules, and generally outperforming baseline models. Specifically, Hot2Mol achieved an average docking score of -8.197 kcal/mol, compared to -7.893 kcal/mol for bioactive molecules, -7.456 kcal/mol for PGMG, and -7.436 kcal/mol for Pocket2Mol, and -7.280 kcal/mol for Lingo3DMol (Supplementary Table S5). Therefore, Hot2Mol is effective in designing PPI inhibitors with relatively strong binding affinities.

**Fig. 4.**
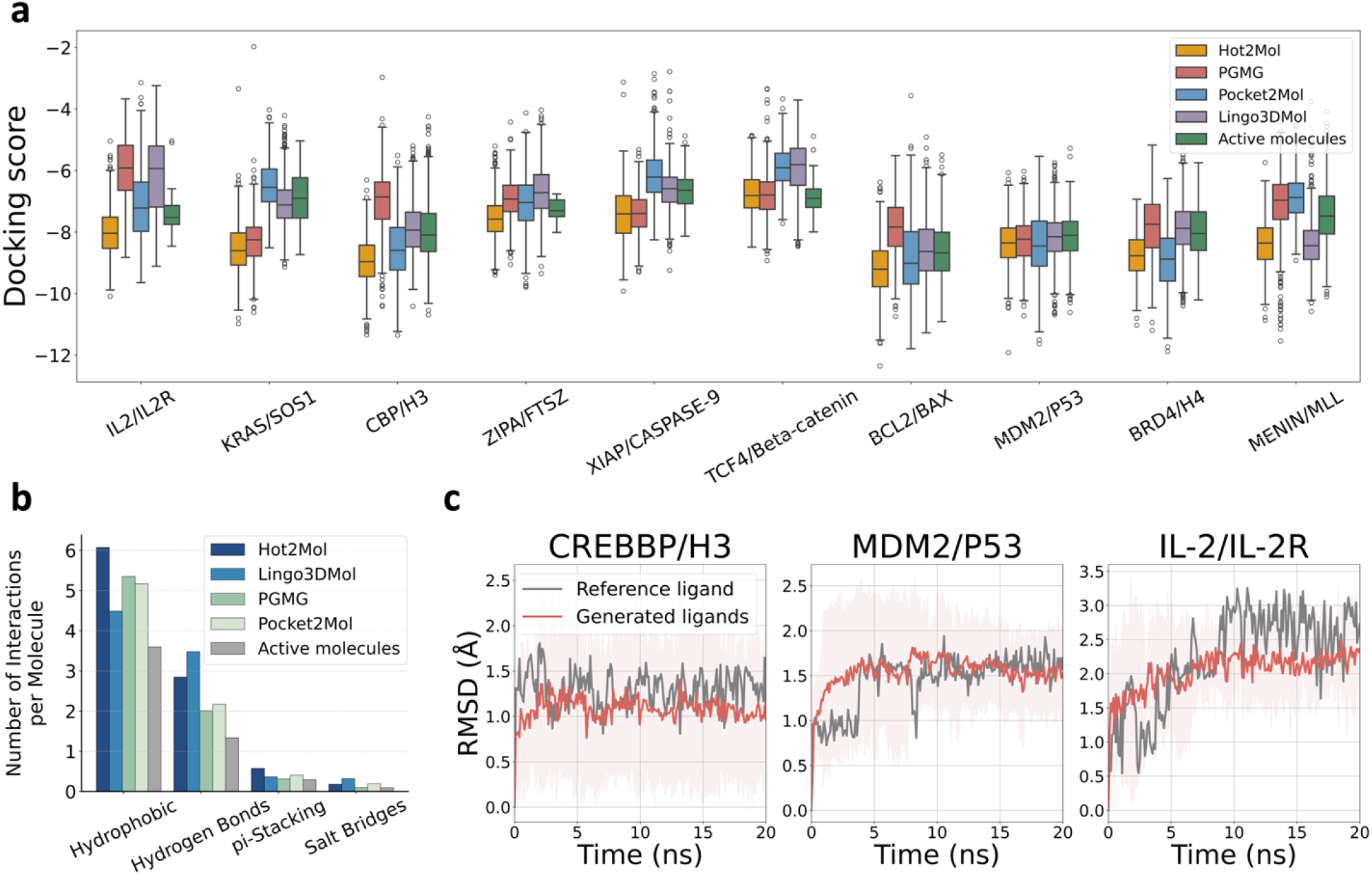
(**a**) Distribution of docking scores for molecules generated by different methods, with each method generating 10,000 molecules across 10 PPI targets. These scores are compared to known bioactive molecules (activity values ≤10 μM) from the DLiP database. The boxplot’s centerline represents the median, with bounds indicating the first and third quartiles. Whiskers extend to 1.5 times the interquartile range (IQR). (**b**) Bar plot showing the number of interactions per molecule for each interaction type, derived from docked poses of molecules generated by different models and known bioactive ligands. (**c**) Ligand RMSD plots during short MD simulations, assessing the binding pose stability of generated ligands against three PPI targets from the test set: MDM2/P53, CBP/H3, and IL-2/IL-2R. Red curves show average RMSDs of five sampled ligands from each generated set with 95% confidence intervals, while gray curves show RMSDs of reference ligands.

To further assess the quality of protein-ligand complexes, we analyzed interaction types using PLIP software ^34^, identifying and quantifying four common intermolecular interactions, which are normalized to determine interactions per molecule (Fig. 4b). These include hydrophobic interactions, hydrogen bonds, π–π stacking, and salt bridges. The greater number of interactions indicate a higher likelihood of strong binding. Hot2Mol-generated molecules consistently showed more interactions per molecule than bioactive molecules, with nearly twice as many hydrogen bonds as the bioactive compounds. Hot2Mol also outperformed Pocket2Mol and PGMG across all interaction types. Although Lingo3DMol generated the highest number of hydrogen bonds among all models, it produced significantly fewer hydrophobic interactions than Hot2Mol. This is likely because Lingo3DMol-generated molecules had lower shape complementarity with target proteins, as indicated by their higher docking scores compared to Hot2Mol.

In Supplementary Fig. S7, we illustrated the distance distributions for the four interaction types, revealing similar distributions across all models. For example, hydrophobic interactions peak around 3.7 Å, while hydrogen bonds peak near 2.25 Å. This consistency indicates that interaction distances are intrinsic to the protein-ligand binding processes, independent of the molecule generation methods. However, the choice of method significantly influences the type and frequency of intermolecular interactions.

### Binding pose stability analysis

To assess the binding stability of Hot2Mol-generated ligands, we conducted 20 ns molecular dynamics (MD) simulations. Ligands that interact unfavorably with their targets often show significant fluctuations in their binding poses over short time periods ^35^. We quantified these fluctuations using root-mean-square deviations (RMSDs) of the ligand poses throughout the MD simulations.

Given the structural diversity of PPI targets, we focused on three representative classes ^36^: (i) globular protein-helical peptide interactions, (ii) globular protein-globular protein interactions, and (iii) globular protein-peptide interactions, with descriptions in Supplementary Fig. S8. A therapeutically relevant target was selected from each class: CBP/H3, MDM2/p53, and IL-2/IL-2R. The same protein structures were used for docking as in previous sections. For each target, 10,000 molecules were generated, and five ligands were randomly sampled from the top 50 molecules with the highest docking scores for MD simulations. For the CBP/H3 interaction, we used the phase 2 small-molecule inhibitor Inobrodib as the reference ligand ^37^. For the MDM2/p53 interaction, the phase 3 small-molecule inhibitor Idasanutlin was chosen as the reference ligand ^38^. Due to the absence of an IL-2/IL-2R inhibitor in clinical trials, we chose a potent inhibitor from the crystal structure 1PY2 as a reference ligand ^39^.

Fig. 4c displays the average RMSD values for the generated ligands (in red) compared to those of the reference ligands (in grey). The RMSD profiles suggest that the generated ligands generally exhibit stable binding, as evidenced by their lower or comparable RMSD values relative to the reference ligands. In the CBP/H3 interaction, the generated ligands demonstrated slightly greater stability than the reference ligand, as reflected by their lower and more consistent RMSD values. For the MDM2/p53 interaction, the generated ligands exhibited similar stability to the reference ligand, with overlapping RMSD profiles. Notably, the generated ligands for the IL-2/IL-2R interaction displayed significantly enhanced stability, suggesting potential for improved therapeutic efficacy. These findings demonstrate that Hot2Mol is capable of generating ligands with both high binding affinity and binding stability for various PPI targets.

### Case studies

#### Structure-based design of MDM2/p53 inhibitors

In this section, we demonstrate the practical utility of Hot2Mol in designing high-affinity inhibitors that target the hot-spot residues involved in the MDM2/p53 interaction. This interaction plays a crucial role in regulating the tumor suppressor p53, with MDM2 inhibiting p53’s activity. Blocking this interaction with small-molecule inhibitors is a promising cancer therapeutic strategy ^40,41^. Idasanutlin (RG7388) is a well-known small-molecule inhibitor of the MDM2/p53 interaction that has progressed into clinical trials for various cancers ^38^.

We used Hot2Mol to generate MDM2/p53 inhibitors by mimicking pharmacophoric features sampled from p53 hot-spot residues Leu26, Trp23, and Phe19, as identified in the crystal structure 1YCR. These molecules were docked against the MDM2 structure (PDB: 3JZK) using the ligand position in the crystal structure as the docking site. An initial library of 10,000 molecules was generated. As shown in Fig. 5a, the QEPPI values of the generated molecules closely align with those of known bioactive molecules, with some optimization observed. The generated molecules exhibited significantly lower SA scores compared to bioactive molecules, indicating enhanced synthetic accessibility. Additionally, these molecules possess a substantially lower molecular weight while still achieving comparable binding affinity (Fig. 4a), indicating better efficacy in disrupting the MDM2/p53 interaction.

**Fig. 5.**
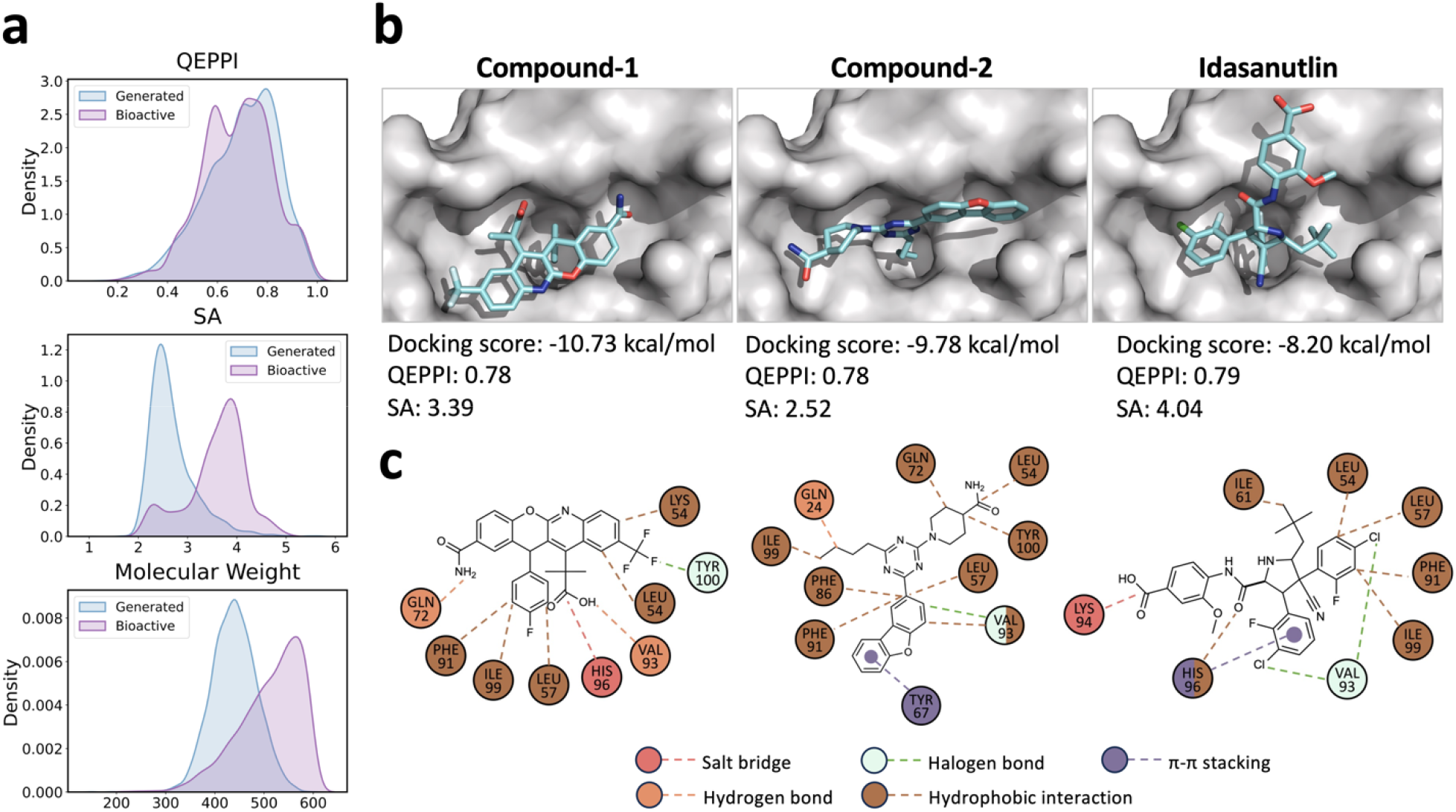
Structure-based drug design targeting the MDM2/p53 interaction. (**a**) Distributions of different molecular properties related to all generated MDM2/p53 inhibitors and known bioactive molecules. Distribution of various molecular properties for the generated MDM2/p53 inhibitors compared to known bioactive molecules. QEPPI represents PPI-targeting drug-likeness, SA is synthetic accessibility, molecular weight is shown in Daltons. (**b**) Docking poses of the generated ligands and Idasanutlin against MDM2 (PDB: 3JZK). (**c**) The 2D interaction diagrams between MDM2 and the generated ligands or Idasanutlin, presented in the same order as in (b). Circles represent amino acid residues, while dashed lines indicate interactions.

The generated molecules were filtered based on (i) QEPPI scores greater than 0.5, (ii) SA scores less than 3.0, and (iii) molecular weight less than or equal to 600, followed by ranking according to docking scores. From the top ten molecules, we selected the two with the best docking poses by visual inspection and compared their binding patterns with Idasanutlin. Visualization in PyMOL ^42^ revealed that the hit compounds effectively bound to MDM2 at sites typically occupied by key P53 residues, namely, Leu26, Trp23, and Phe19 (Fig. 5b). In addition to achieving lower docking scores than Idasanutlin, the generated ligands displayed similar QEPPI values and significantly lower SA scores, indicating that these compounds not only have a strong potential for effective binding but are also more synthetically accessible and drug-like, making them promising candidates for further optimization. Fig. 5c shows the 2D interaction maps of the three molecules with MDMD2, generated using PLIP software ^34^. The generated ligands interacted with key residues on MDM2 through various interactions, including hydrogen bonds, salt bridges, and halogen bonds, with residues including Val93, Leu54, Leu57, Phe91, and Ile99, similar to Idasanutlin. Notably, the generated ligands engaged additional hot-spot residues compared to Idasanutlin, including Tyr100 and Gln72.

In addition to the MDM2/p53 target, we also demonstrated structure-based design of inhibitors for the IL-2/IL-2R and CBP/H3 interactions (Supplementary Analyses). These results demonstrate Hot2Mol’s capability to generalize across diverse PPI structures.

#### Selectivity-controlled design of BCL-XL/BAK inhibitors

In this section, we address a practical challenge where inhibitor selectivity is crucial. BCL-XL is a member of the anti-apoptotic BCL2 protein family, which regulates cell death by inhibiting pro-apoptotic proteins like BAK ^43^. Overexpression of BCL-XL is linked to cancer cell resistance to apoptosis, hence making it a critical target for selective inhibitors. Achieving selectivity for BCL-XL over other BCL2 family members is essential to minimize side effects and enhance therapeutic efficacy. Lessene et al. ^44^ designed a selective inhibitor for the BCL-XL/BAK interaction, WEHI-539 (PDB: 3ZLR), which demonstrated more than 500-fold higher affinity for BCL-XL compared to BCL2. Therefore, we chose WEHI-539 as the reference ligand.

We initially generated 10,000 molecules targeting the BCL-XL/BAK interaction using Hot2Mol. Filtering for synthetic accessibility (SA) scores below 3, QEPPI scores above 0.5, and molecular weights under 600 yielded 6,567 molecules. The docking scores of the generated ligands for both BCL-XL and BCL2 are shown in Fig. 6a. Points above the solid diagonal line indicate lower scores for BCL-XL compared to BCL-2, suggesting stronger binding to BCL-XL. Our results show that the majority of generated ligands bind more strongly to BCL-XL, as evidenced by the higher concentration of samples above the solid line. A reduction in energy by 1.36 kcal/mol theoretically corresponds to a 10-fold decrease in inhibitory concentration. Therefore, we set a criterion for selectivity as a 2.72 kcal/mol difference in docking scores between BCL-XL and BCL2, equivalent to a 100-fold difference in inhibitory concentration ^21^. This filtering yielded 74 selective ligands, as highlighted by the red points above the dashed line. We selected one ligand with the best docking pose based on visual inspection (Fig. 6b).

**Fig. 6.**
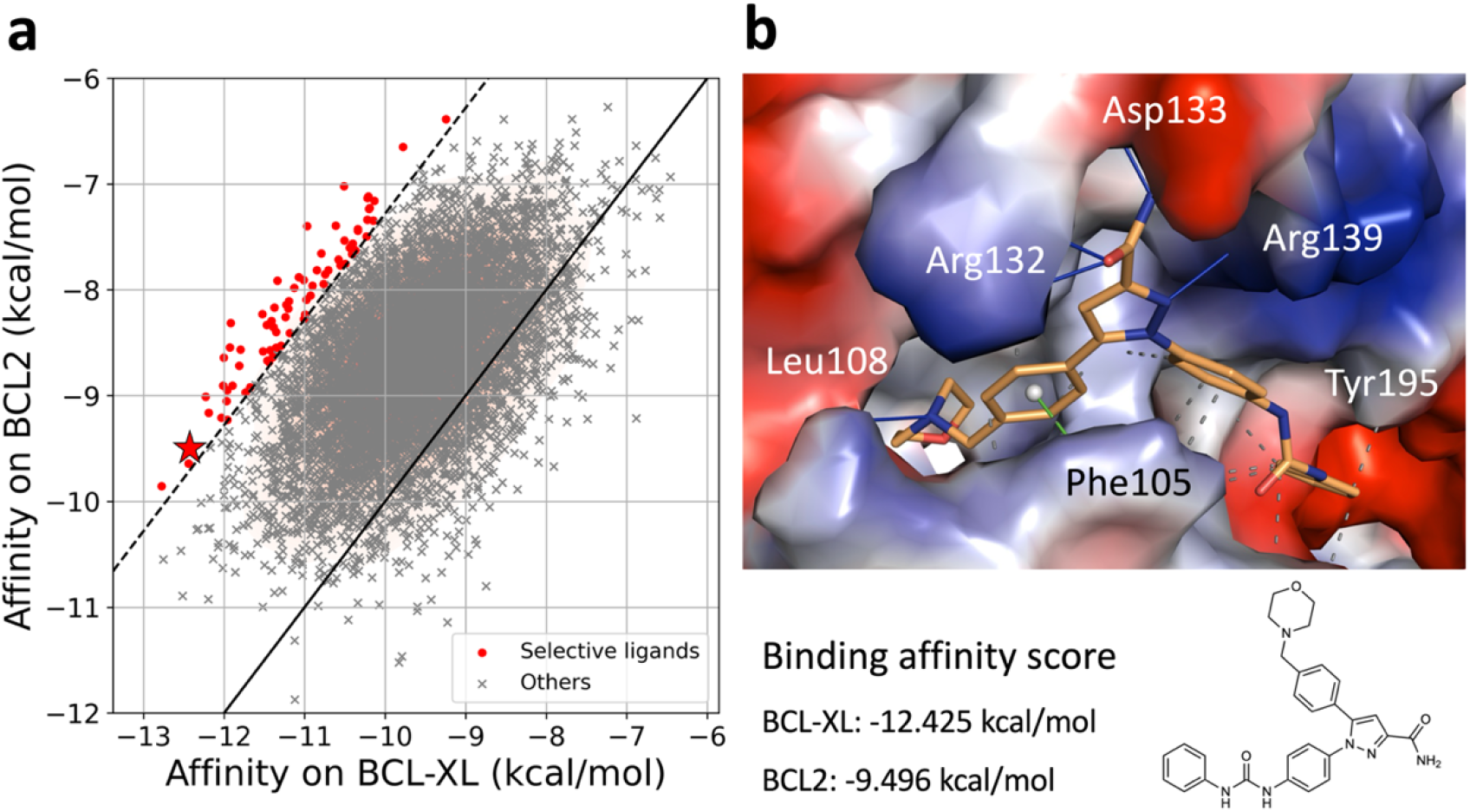
Selectivity-controlled PPI drug development scenario. (a) Scatter plot showing the binding affinity scores of the generated BCL-XL/BAK inhibitors against BCL-XL and BCL2 proteins. Red points indicate a 2.72 kcal/mol higher binding affinity for BCL-XL, corresponding to a 100-fold lower inhibitory concentration. (b) Docked pose of a well-designed ligand (highlighted with a star in (a)) expected to be selective for BCL-XL (PDB: 3ZLR). Non-covalent interactions with key binding residues are shown: blue lines for hydrogen bonds, green dashed line for *π*-*π* interactions, and grey dashed lines for hydrophobic interactions.

The hit ligand binds within the hydrophobic groove on BCL-XL, engaging key residues in pockets P2 and P4, which are critical for selective binding of BCL-XL inhibitors ^44^. The strongest interactions include hydrogen bonds with Arg132, Arg139, Asp133, and Leu108; π–π interaction with Phe105; and hydrophobic interactions with Tyr195 and Phe105. The generated ligand also aligns well with WEHI-539 (Supplementary Fig. S9), indicating a good fit within the binding groove. This is reflected in the ligand’s binding affinity for BCL-XL (−12.425 kcal/mol), comparable to that of WEHI-539 (−12.6 kcal/mol). In contrast, the ligand shows a much lower binding affinity for BCL2 (−9.496 kcal/mol vs -10.5 kcal/mol), indicating improved selectivity. In summary, we have identified a promising molecule with high affinity and selectivity for BCL-XL, making it a strong candidate for targeted therapy.

### Ablation studies

We conducted ablation analyses to evaluate the contribution of each key component to the superior performance of Hot2Mol, including the equivariance constraint in geometric space and the imposition of molecular property constraints. We developed two modified models: the without-EGNN model and the without -property-constraint model. The without-EGNN model replaces the EGNN encoders with a Gated Graph Convolutional Network ^45^, employs a 2D topology graph in place of a 3D molecular graph, and substitutes the 3D pharmacophore graph with a fully connected 2D graph lacking spatial information. The without-property-constraint model removes the encoding for molecular property constraint. Both models were tested on the same 10 PPI targets, generating 10,000 molecules each.

As shown in Table 2 and Supplementary table S6, omitting the spatial information of molecules and pharmacophores significantly reduces the fraction of valid molecules generated (from 0.971 to 0.904). Additionally, the model generates more repeated molecules (uniqueness drops from 0.973 to 0.67), indicating reduced exploration of chemical space during training. This reduction in the validity and uniqueness underscores the importance of spatial information and equivariance constraint in geometric space. The SA score for molecules from the without-EGNN model is 2.868, suggesting that these molecules are generally more difficult to synthesize compared to those from the full model (SA=2.692). Furthermore, the mean docking score increased from -8.229 kcal/mol to -7.848 kcal/mol after removing the EGNN module, indicating weaker binding affinity of the generated molecules to targets.

**Table 2.** Results of the ablation analysis.

|  | Validity<br>↑ | Uniqueness↑ | Novelty↑ | Ratio of<br>Available<br>Molecules↑ | SA↓ | QEPPi↑ | Mean<br>Docking<br>Score↑ |
| --- | --- | --- | --- | --- | --- | --- | --- |
| Original | **0.971** | 0.973 | 0.999 | **0.943** | **2.692** | **0.654** | -8.229 |
| without-EGNN | 0.904 | 0.67 | **1.0** | 0.617 | 2.868 | 0.496 | -7.848 |
| without-property-constraint | 0.936 | **0.995** | **1.0** | 0.931 | 2.902 | 0.506 | **-8.385** |

An upward arrow indicates that higher values represent better performance, and vice versa. The best-performing variant for each metric is highlighted in **bold**.

The without-property-constraint model generates a similar ratio of available molecules and docking scores as the full model. However, the average QEPPI scores drop from 0.654 to 0.506, demonstrating the effectiveness of the property constraint in optimizing drug-like properties. Additionally, the generated molecules are harder to synthesize, with the SA score increasing from 2.692 to 2.902. In summary, the inclusion of spatial information in the molecular structure and pharmacophore hypotheses, along with the use of EGNNs for embedding, is crucial for generating valid and high-affinity PPI modulators. Furthermore, the QEPPI property constraint is crucial for ensuring that the generated molecules have good druggability.

## Discussion

In this study, we introduce Hot2Mol, a novel deep generative framework for the structure-based design of PPI inhibitors, specifically addressing the complexities of PPI interfaces. Unlike conventional drug targets with well-defined cavities or pockets, PPI interfaces are typically flatter and more hydrophobic, posing challenges for small-molecule binding. To overcome these challenges, Hot2Mol integrates deep molecular generative modeling with prior knowledge of PPI targets, utilizing pharmacophores derived from hot-spot residues to guide the generation of small molecules. By directly mimicking the pharmacophores of hot-spot residues, Hot2Mol allows for precise and efficient targeting of key regions, leading to the generation of high-affinity and selective inhibitors. Additionally, by targeting key interaction sites and using a pharmacophore-guided design approach, Hot2Mol improves the interpretability of DGMs, offering clearer insights into the mechanisms of PPI inhibition.

Hot2Mol utilizes EGNNs to accurately encode the 3D molecular structures and the spatial distribution of pharmacophores, keeping equivariance properties in geometric space. A conditional transformer is employed to model the relationships between molecular structures, pharmacophore features, and molecular properties, guiding the generation of small molecules that are simultaneously optimized for binding affinity and drug-like characteristics. The introduction of latent variables allows broader exploration of the chemical space, enhancing the diversity and novelty of the generated PPI modulators. Notably, Hot2Mol is flexible and highly generalizable, as it does not rely on bioactive ligands for fine-tuning. This approach avoids the common issue of ligand scarcity in PPI drug development.

Our experiments show that Hot2Mol generates PPI inhibitors with superior validity, uniqueness, drug-likeness, and synthetic accessibility compared to state-of-the-art models. These inhibitors achieve binding affinities that match or exceed those of known bioactive compounds while outperforming baseline models. Chemical space analysis confirms that the generated molecules accurately capture natural distributions and maintain geometric validity. Additionally, Hot2Mol-generated molecules form more interactions with target proteins than bioactive compounds, and MD simulations confirm their binding stability across diverse PPI structures.

The case studies further illustrate Hot2Mol’s practical utility in designing PPI inhibitors that effectively target hot-spot residues. This capability is demonstrated across a diverse set of PPI targets, including MDM2/p53 (globular protein-helical peptide interaction), IL-2/IL-2R (globular protein-globular protein interaction), and CBP/H3 (globular protein-peptide interaction). Additionally, we show that Hot2Mol can generate highly selective inhibitors, as exemplified by the BCL-XL/BAK interaction.

Overall, Hot2Mol demonstrates significant potential to accelerate PPI drug discovery by leveraging AI techniques and deep structural insights into PPI targets. However, there is still room for improvement. For instance, integrating protein-ligand interaction rules may further enhance the binding affinities of generated molecules ^21^. Additionally, simultaneous optimization of multiple molecular properties is another area for future research.

## Methods

### Data preparation

To train the Hot2Mol model, a dataset comprising Matched Molecular Pairs (MMPs) was used, which were extracted from ChEMBL 28 ^46^, following the methodology in previous research ^47^. MMPs are pairs of molecules that differ by a single transformation, a common strategy in medicinal chemistry for molecular optimization. The MMPs were constructed using the open-source tool mmpdb ^48^.

MMPs were selected from molecules reported within the same publication to increase the likelihood of shared binding pockets, thereby more accurately reflecting chemists’ intuition and practical considerations. The data preparation involved several key steps, as outlined below:

#### Molecular pair pre-processing

⍰ Standardization using MolVS ^49^
⍰ Molecules with 10 to 50 heavy atoms
⍰ Number of rings > 0
⍰ AZFilter = “CORE” ^50^ to exclude low-quality compounds
⍰ Use substructure filters ^51^ for hit triaging, SeverityScore < 10 ^52^

#### Publication pre-processing

⍰ Publications from the year 2000 or later
⍰ Publications containing between 10 and 60 molecules

#### Further molecular pair pre-processing

⍰ Remove duplicated pairs
⍰ Exclude reverse pairs

Following these steps, 320,563 pairs were randomly sampled for training, and 21,362 pairs were randomly sampled for validation.

The structure of the PPI partner proteins used for docking analysis are MENIN/MLL (PDB: 6WNH), IL-2/IL-2R (PDB: 1PY2), KRAS/SOS1 (PDB: 7UKR), CBP/H3 (PDB: 5W0F), ZIPA/FTSZ (PDB: 1S1J), XIAP/CASPASE-9 (PDB: 1TFT), BCL2/BAX (PDB: 6O0K), MDM2/P53 (PDB: 3JZK), TCF4/Beta-catenin (PDB: 7ZRB), and BRD4/H4 (PDB: 3P5O).

### Details of Hot2Mol

Hot2Mol is a molecular generative model designed for de novo generation of target-specific and drug-like PPI inhibitors. It utilizes a structure-based design approach by mimicking the pharmacophores of hot-spot residues at PPI interfaces, therefore achieving competitive inhibition of PPI target proteins. This methodology is framed as a mapping learning problem, where the model is trained to capture the many-to-many relationships between molecular structures and conditions. Specifically, the model learns the mapping of pharmacophoric features ***A***, molecular property conditions ***B***, and molecular structures ***C*** as (***A, B***) **→ *C***. During the generation process, the model samples from a base distribution to autoregressively generate molecules that align with the hot-spot pharmacophore distributions and are optimized for drug-like properties. The architecture of Hot2Mol is based on a variational autoencoder (VAE) that consists of an encoder and a decoder (Fig. 1).

The model employs a conditional transformer encoder to map the input embeddings into a shared latent space using a latent vector *z*. The inputs include a 3D molecular graph ***X***, a 3D graph representation of pharmacophore hypothesis ***P***, and molecular property encoding ***D*** (Fig. 1c). This configuration enables the model to learn a 3D-aware latent space, capturing the relationships between a molecule’s 3D structure, its pharmacophore distributions, and drug-like property constraints. To ensure equivariance under rotations, translations, reflections, and permutations in geometric space, E(n)-equivariant neural networks (EGNNs) ^53^ are utilized to embed both the molecular graph ***X*** and the pharmacophore graph ***P***. Additionally, a Multi-Layer Perceptron (MLP) is used to embed the one-hot property encoding ***D*** into a dense feature vector (Fig. 1e). These embeddings are concatenated and input to the transformer encoder:

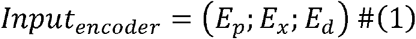

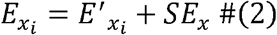

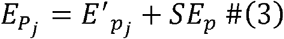

Here, *E*_*d*_ is the embedding vector of the molecular property constraint, 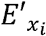 is the graph embedding vector of the z-th atom, 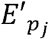. is the graph embedding vector of the *j-*th pharmacophore feature, and *SE*_*X*_ and *SE*_*p*_ are the segment embedding vectors for the molecule and pharmacophore hypothesis, respectively. After several layers of the transformer encoder block, an attention pooling layer averages the combined features to derive the final molecule representation *h*_*x*_. This representation *h*_*x*_ is then processed by two separate subnetworks to compute the mean *μ* and the log variance *logσ*^*2*^ of the posterior variational approximation. Latent variables *z* are subsequently drawn from a normal distribution *N*(*μ*, Σ).

The transformer decoder autoregressively generates molecules in SMILES format from the latent space, integrating the pharmacophore and molecular property embeddings as conditions (Fig. 1d). The decoder network takes the latent variables, pharmacophore embeddings, and property embeddings as inputs:

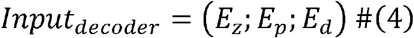

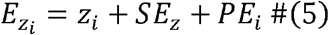

In this context, *E*_*d*_ represents the embeddings of the property constraints, *SE*_*Z*_ denotes the segment embeddings for the latent variables, and *PE*_*i*_ represents the positional embeddings for the *i*-th token. The decoder generates the SMILES of the target molecule in an autoregressive manner, with each token being predicted based on the previously generated tokens and the conditional inputs:

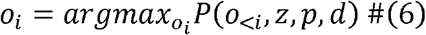

where o_*i*_ is *i*-th generated token. Notably, the decoder shares embedding layers with the encoder for pharmacophores and property conditions.

### Representation of molecules

Molecules are represented as a 3D molecular graph *G*_*x*_ = (*x,r,e*). In this representation, *x* = (*x*_*1*_, *x*_*2*_, …, *x*_*n*_) ∈{0,l}^*n*×*f*^ denotes the one-hot encoded atom types, and *r* = (*r*_1_..*r*_2_., *r*_*n*_) ∈ *R*^*n*×3^ represents the continuous atom coordinates in 3D space. The edge attributes *e* = (*e*_*ij*_) indicates the existence of covalent bonds between atoms. The 3D molecular conformer is constructed using RDKit ^31^ based on SMILES notation.

### Representation of pharmacophores

To generate molecules that align with specific pharmacophore distributions, we represent pharmacophore hypothesis using a fully connected 3D graph, denoted as *G*_*p*_ = (*f,p,e*), where *f* = *f*_1_, *f*_2_, … *f*_*m*_) ∈ {0,l}^*m*×*k*^ represents the one-hot encoded feature types (such as aromatic, hydrogen acceptor, hydrogen donor, etc.), and *p* = (*p*_*1*_, *p*_*2*_ > …, *p*_*m*_) ∈ *R*^*m*×3^ represents the continuous 3D coordinates of each pharmacophore feature, and *e* = (*e*_*ij*_) represents the edges in the graph, indicating the distances between pairs of pharmacophores.

During the training stage, pharmacophore graphs are constructed from small molecules in the training set (Fig. 1f). The pharmacophoric features of these molecules are computed using RDKit, and a random subset of features is sampled for each molecule. This approach enables the model to learn the general relationship between pharmacophore patterns and molecular structures, facilitating the generation of molecules that fit specific pharmacophore hypotheses.

For the inference stage, the generation process is guided by the pharmacophoric features derived from the hot-spot residues of the PPI target (Fig. 1f). To identify these hot-spot residues, we employed HawkDock ^54^, which combines the ATTRACT docking algorithm with MM/GBSA (Molecular Mechanics/Generalized Born Surface Area) free energy decomposition analysis. MM/GBSA predicts binding free energies and decomposes these contributions on a per-residue basis. The top three residues with the lowest binding energies on the PPI target protein are designated as the hot-spot residues. Pharmacophoric features are then computed based on these hot-spot residues, and a subset is randomly selected to serve as the pharmacophore hypothesis. This pharmacophore hypothesis guides the generation of molecules, ensuring they mimic the intermolecular interaction modes of the hot-spot regions.

### Encoding of molecular property constraints

To generate PPI inhibitors with favorable drug-like properties, constraints are imposed during molecule generation using the Quantitative Estimate of PPI-targeting drug-likeness (QEPPI) ^55^. QEPPI is an extension of the Quantitative Estimate of Drug-likeness (QED) method^56^.

Property changes between source molecules and their corresponding target molecules are calculated by measuring the difference in their QEPPI scores. This allows the model to learn the relationship between specific molecular transformations and their impact on QEPPI. Matched molecular pairs {***A, B, C***} are generated, where ***A*** is the source molecule, ***B*** is the target molecule, and ***C*** is the QEPPI property change from ***A*** to ***B***. The model is trained to map the input space (*A, C*) ∈ *A* × *C* to the output space *B* ∈ *B*. Property changes, as continuous values, are divided into intervals with a width of 0.1 units. These intervals are encoded as one-hot vectors ***D***, which are then processed through a linear neural network layer to generate embeddings *E*_*d*_.

During the molecule generation process, all negative QEPPI change tokens are masked, and positive change tokens are set to 1, ensuring that the generated molecules are optimized for QEPPI score. This approach ensures that generated PPI inhibitors exhibit favorable drug-like properties, making them suitable candidates for further drug development.

### Loss function

The loss function for Hot2Mol incorporates three components: KL divergence loss, sequence reconstruction loss, and pharmacophore matching loss:

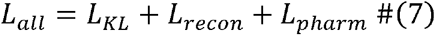

The first two terms are derived from the evidence lower bound (ELBO) of the log-likelihood

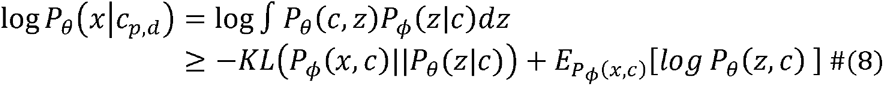

Here, KL represents the Kullback–Leibler divergence, and *P*_*θ*_ (*z*|*c*) is assumed to be a standard Gaussian distribution *N*(*0,1*). The term *KL*(*P*_*ϕ*_ (*x,c*)||*P*_*θ*_ (*z*|*c*)) is the KL loss *L*_*KL*_, which serves as a regularization term to align the prior distribution with the posterior, thus smoothing the latent space of *z*. The expectation term, 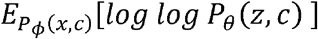, is estimated using Monte Carlo estimation with a single data point per sample ^57^. This term represents the SMILES sequence loss *L*_*recon*_, which is responsible for reconstructing the target sequence.

The pharmacophore matching loss *L*_*pharm*_ assesses the model’s ability to learn the correspondence between heavy atoms and pharmacophore elements. The matching score for the *i*-th pharmacophore *p*_*i*_ and the j-th output token *o*_*j*_ is calculated as follows:

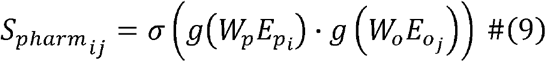

In this equation, 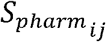 represents the matching score,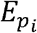. and 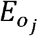. are the embedding vectors for pharmacophores and outputs, respectively. *W*_*p*_ and *W*_*o*_ are learnable matrices that project the two different embeddings into a common space. The dot product is denoted by □, *σ* is the sigmoid function, and *g* is the ReLU function. The pharmacophore matching loss *L*_*pharm*_ is then computed as the cross-entropy between the matching scores *S*_*pharm*_ and the true labels.

### Training

The EGNN encoders for both pharmacophore graph and molecular graph consist of 4 layers with a hidden dimension of 384. The Transformer encoder and decoder each have 8 layers with 8-head attention mechanisms, 384 hidden dimensions, and 1024 feed-forward dimensions. The model was trained using the Adam optimizer with a learning rate of 3e-4 and a weight decay rate of 1e-6, over 32 epochs. To prevent posterior collapse, cosine learning rate annealing with a cycle length of four epochs was implemented. Gradient clipping was applied to manage training stability, setting the maximum gradient norm to 5. All training processes were executed on an NVIDIA 3090 GPU.

### Evaluation

To assess the capability of different models to generate novel molecules, seven different metrics are utilized: validity, uniqueness, novelty, the ratio of available molecules, synthetic accessibility (SA) score ^55^, QEPPI, and docking scores. Validity calculates the proportion of chemically valid molecules among all generated SMILES strings. Uniqueness measures the proportion of valid and unique molecules. Novelty indicates the percentage of chemically valid molecules out of the total generated molecules. The ratio of available molecules represents the percentage of novel molecules among all the generated molecules. SA score evaluates the ease of synthesizing the generated molecules, with lower scores indicating higher synthetic accessibility. QEPPI assesses the drug-likeness of compounds targeting PPIs. The docking score from AutoDock Vina ^33^ approximates the binding affinity of the generated molecules to PPI partner proteins. AutoDock Vina performs semiflexible docking, considering ligand flexibility while docking into a rigid receptor, using default parameters. The central coordinates of the docking box are the average coordinates of each heavy atom in the reference ligand, with a globally defined cubic box size of width 30 Å to accommodate potentially larger molecules than the reference ligands.

For evaluating baseline models, including Pocket2Mol ^23^, Lingo3DMol ^26^, and PGMG ^27^, experiments were conducted using their published source code and parameters. For Pocket2Mol and Lingo3DMol, a minor modification was made to the code to prevent filtering out duplicated results, allowing all generated molecules to be output. Additionally, for Lingo3DMol, the pocket structure was extracted as the residues within 5 Å around the ligand in the reference protein-ligand complexes. For PGMG, pharmacophores were randomly sampled from the reference ligand for each PPI target to build the pharmacophore hypotheses.

### Molecular dynamics simulation

During the MD simulations, topology and parameter files for the ligands were generated using the GAFF-2.11 force field ^58^ with Amber tools ^59^. The protein-ligand complexes were prepared using Amber tools’ tleap module. The system was solvated in a cubic box with TIP3P water, extending 10 Å from the protein to ensure adequate padding. It was neutralized by adding *Na*^+^ and *Cl*^−^ ions. The systems were simulated under periodic boundary conditions in the NPT ensemble with a Langevin thermostat. Amber FF14SB force field ^60^ was applied to model the interactions. The simulations used the OpenMM toolkit ^61^, starting with energy minimization and followed by a 1 ns equilibration phase. Production simulations were run at 303.5 K and 1 bar pressure with a 2 fs time step. Each protein-ligand complex underwent a 10 ns production phase. To evaluate ligand binding stability, protein backbone structures from each frame were aligned based on heavy atom coordinates using MDtraj software ^62^. This approach allowed for isolation of ligand movement and computation of its RMSD.

## Data availability

The original protein-protein complex data and protein-ligand complex data used in this study is available in the PDBbind database http://www.pdbbind.org.cn. The processed data for training Hot2Mol are available at https://github.com/sun-heqi/Hot2Mol. Source data files relevant to each figure are provided with this paper.

## Code availability

The implementation of our framework, covering the training and sampling of Hot2Mol is available at https://github.com/sun-heqi/Hot2Mol.

## Acknowledgements

Dong-Qing Wei is supported by grants from the Intergovernmental International Scientific and Technological Innovation and Cooperation Program of The National Key R&D Program(2023YFE0199200), the National Science Foundation of China (Grant No. 32070662 and 32030063). The computations were partially performed at the Pengcheng Lab. and the Center for High-Performance Computing, Shanghai Jiao Tong University.

## Author contributions

H.S. conceived the original ideas for this study, designed and performed the experiments, and wrote the manuscript. J.L., Y.Z., S.L., J.C., H.T. contributed to the model design and revision of the manuscript. R.W, X.M, J.Z. and R.L. helped to prepare figures. Y.X. and D.-Q. W. supervised the work.

## Competing interests

The authors declare no competing interests.

## Notes

### Competing Interest Statement

The authors have declared no competing interest.

