## Supplementary Information for "Target-specific design of drug-like PPI inhibitors via hotspot-guided generative deep learning"

**Supplementary Figures**

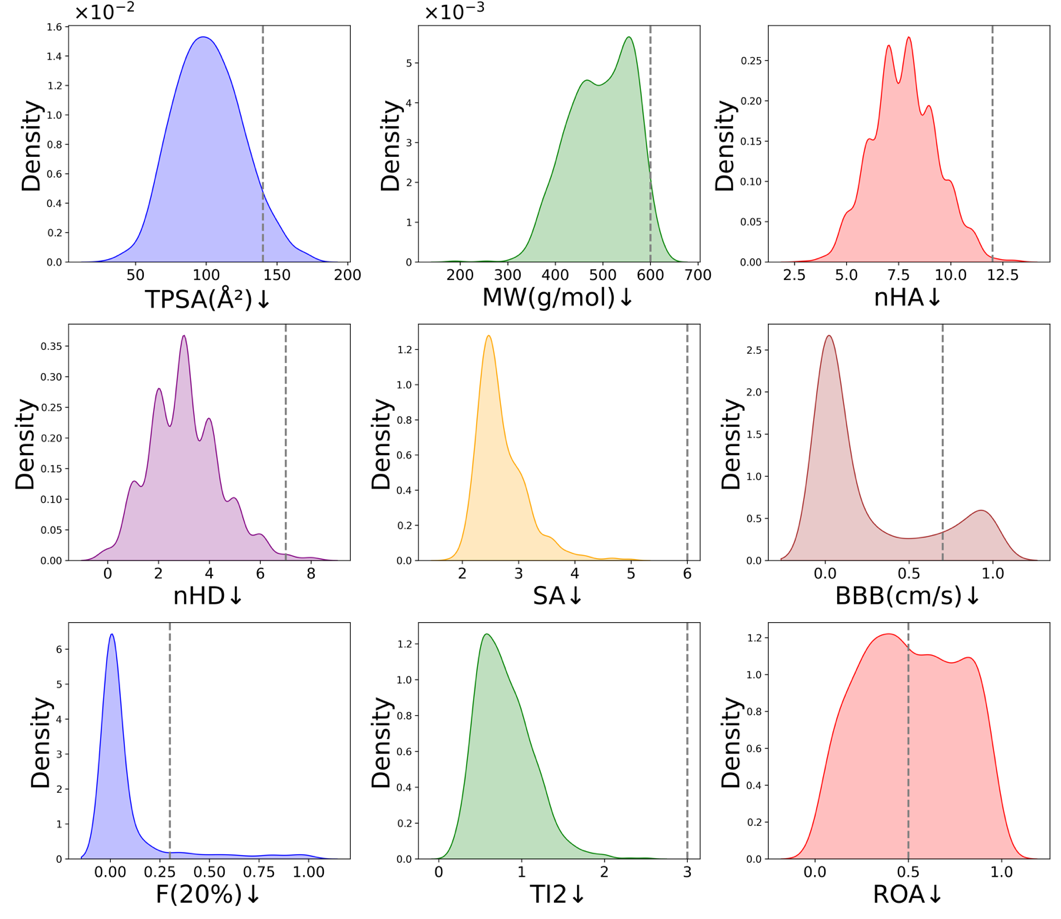

Fig. S1. Distribution of ADMET properties for molecules generated by Hot2Mol. Dashed lines indicate property thresholds: upward arrows signify preferred values above the threshold, and downward arrows signify preferred values below the threshold. TPSA (topological polar surface area): optimal range 0–140 Å²; MW (molecular weight): optimal range 100–600; nHA (number of hydrogen bond acceptors): optimal range 0–12; nHD (number of hydrogen bond donors): optimal range 0–7; SA (synthetic accessibility score): optimal <6; BBB (blood-brain barrier penetration probability): optimal 0–0.7; F(20%) (human oral bioavailability <20% probability): optimal <0.3; T12 (half-life prediction probability ≤3): optimal lower values; ROA (acute toxicity in mammals prediction probability): optimal lower values.

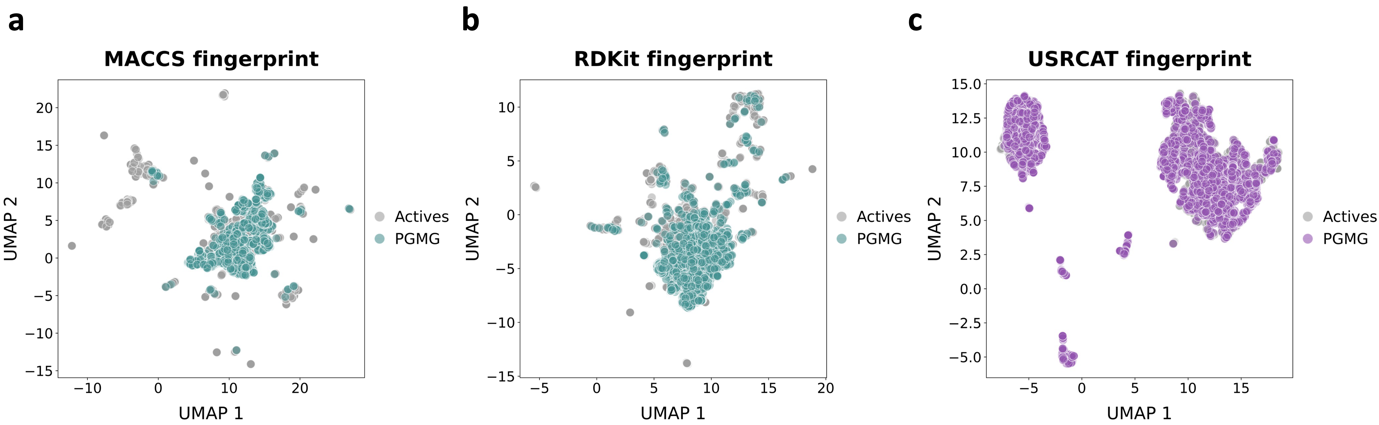

Fig. S2: Visualization of chemical space distribution of PGMG-generated molecules and bioactive molecules. **(a)** MACCS fingerprints, **(b)** RDKit fingerprints, and **(c)** USRCAT fingerprints visualized using UMAP in two-dimensional space.

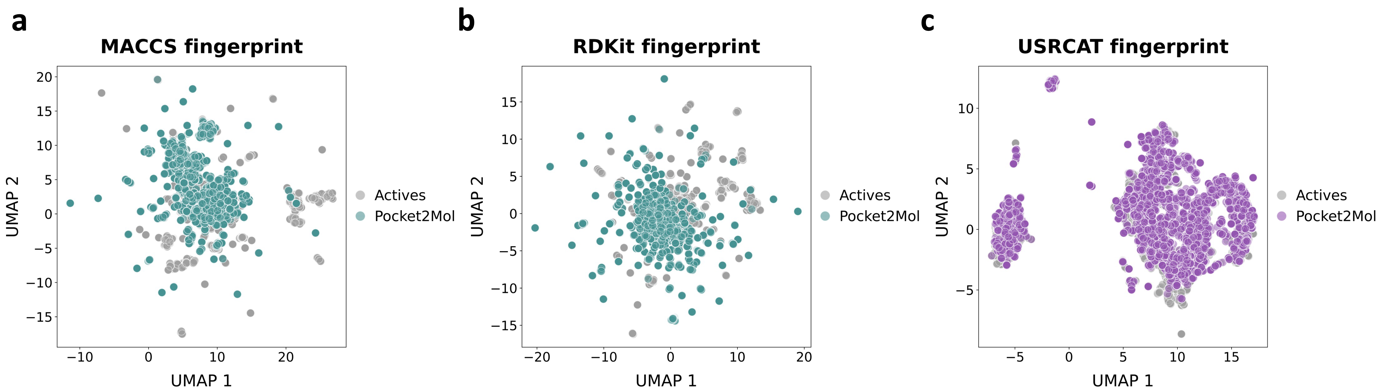

Fig. S3: Visualization of chemical space distribution of Pocket2Mol-generated molecules and bioactive molecules. **(a)** MACCS fingerprints, **(b)** RDKit fingerprints, and **(c)** USRCAT fingerprints visualized using UMAP in two-dimensional space.

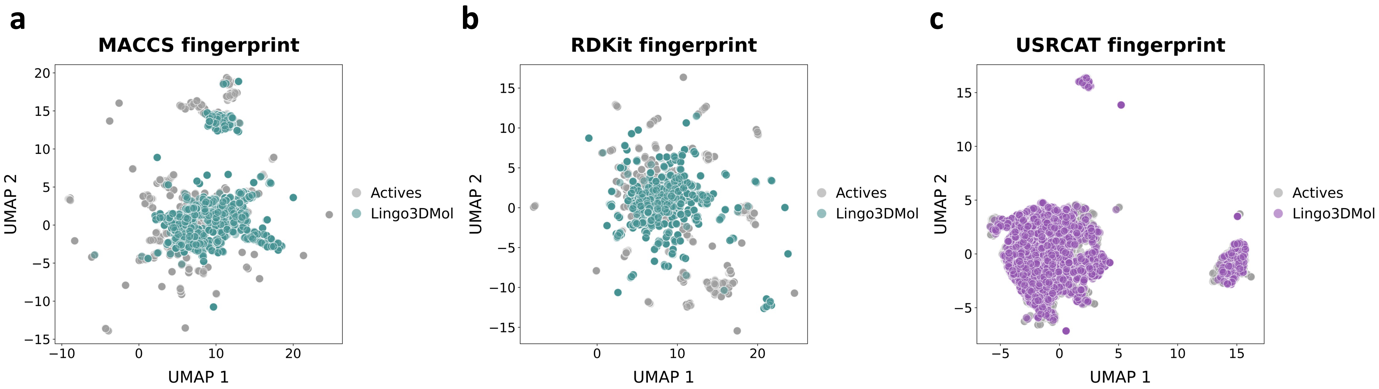

Fig. S4: Visualization of chemical space distribution of Lingo3DMol-generated molecules and bioactive molecules. **(a)** MACCS fingerprints, **(b)** RDKit fingerprints, and **(c)** USRCAT fingerprints visualized using UMAP in two-dimensional space.

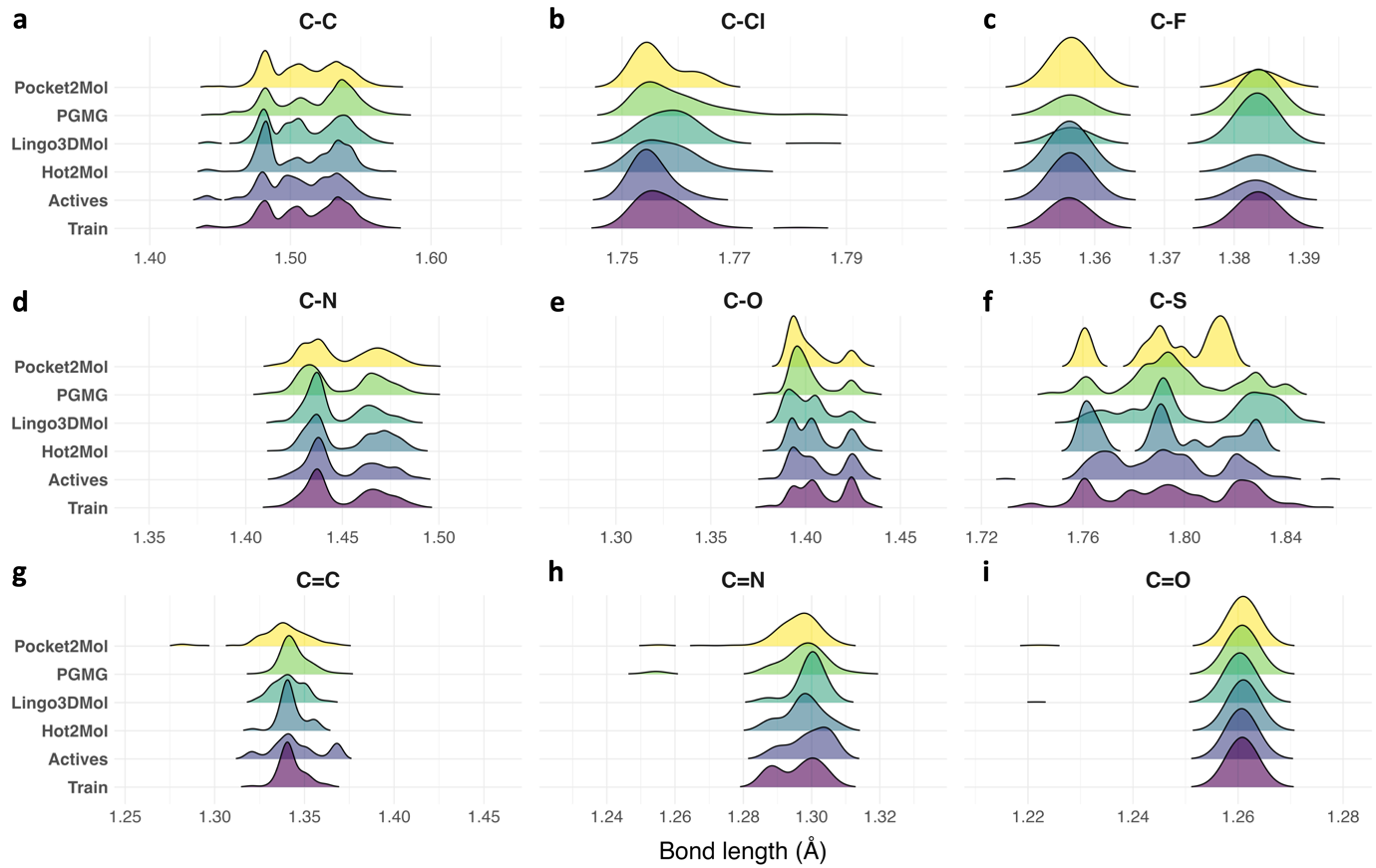
 Fig. S5. Bond length distributions of molecules generated by four DGMs, molecules in the training dataset, and bioactive molecules. The analysis includes nine types of chemical bonds: (a) C-C, (b) C-Cl, (c) C-F, (d) C-N, (e), C-O, (f) C-S, (g) C=C, (h) C=N, and (i) C=O.

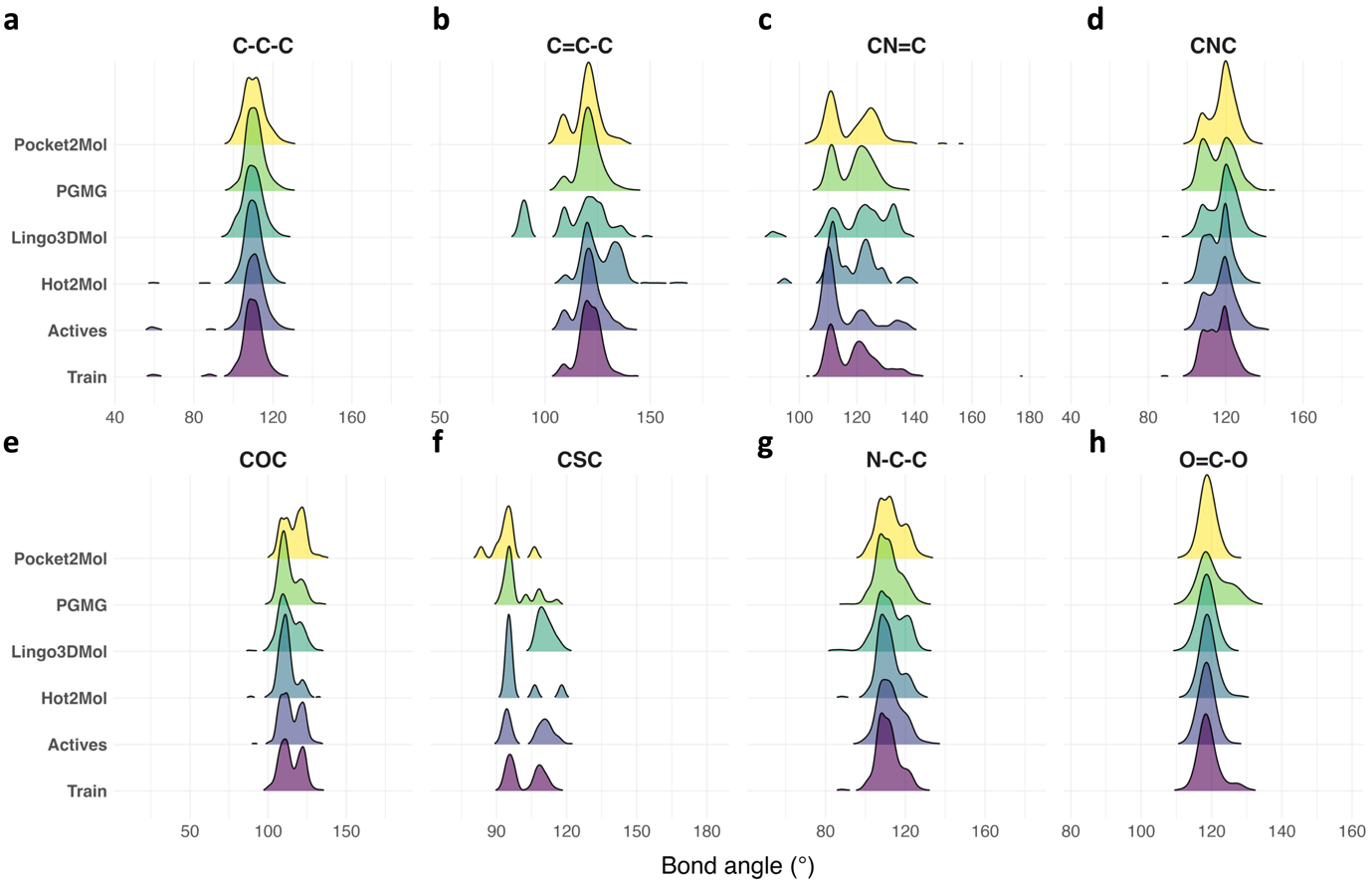

Fig. S6. Bond angle distributions of molecules generated by four DGMs, molecules in the training dataset, and bioactive molecules. The analysis includes eight types of bond angles: (a) CCC, (b) C=CC, (c) CN=C, (d) CNC, (e) COC, (f) CSC, (g) NCC, (h) O=CO.

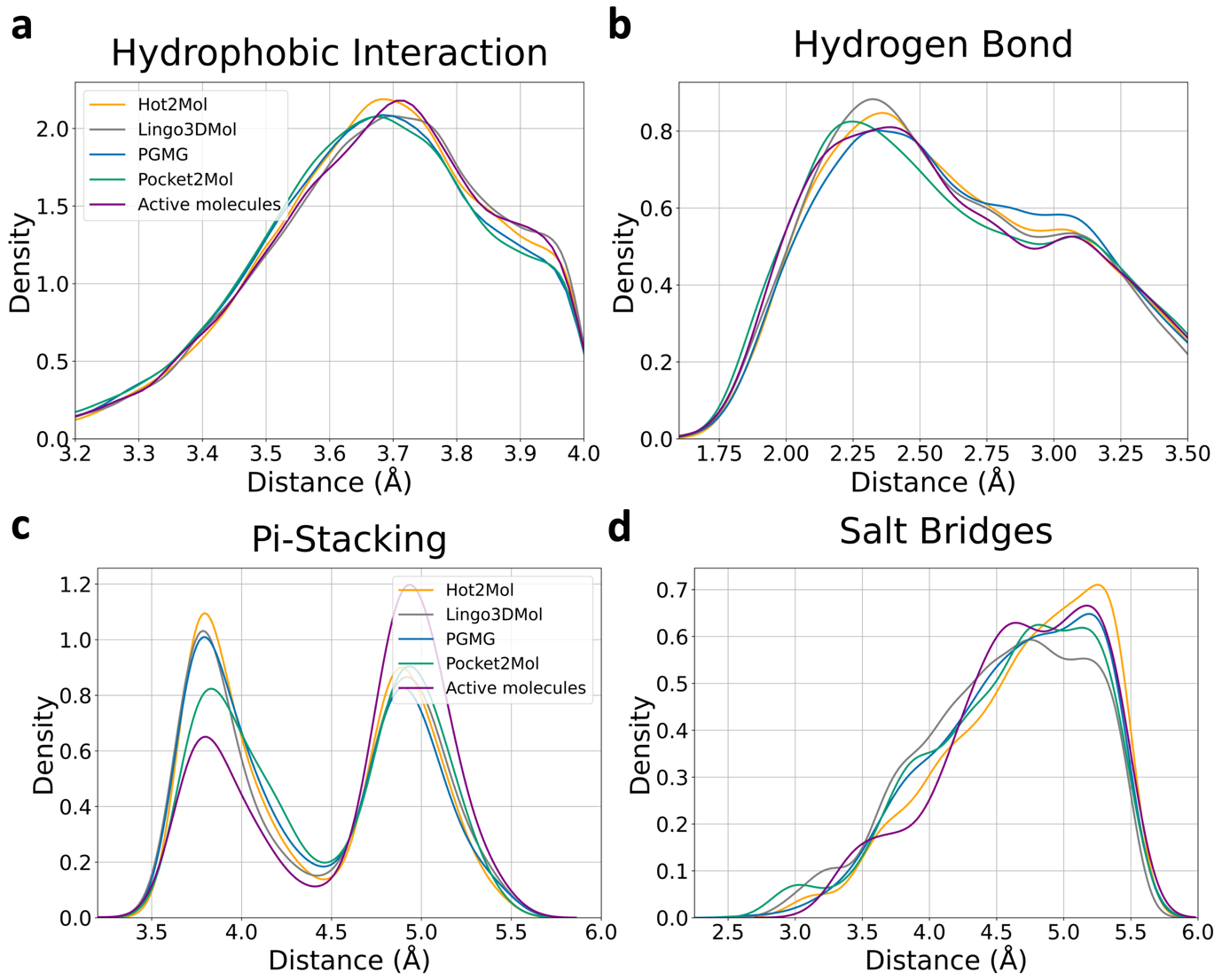

Fig. S7. Distribution of interaction distances in generated and experimentally validated active complexes of ligands and targets. The interactions include: (a) hydrophobic interactions, (b) hydrogen bonds, (c) π-π stacking, and (d) salt bridges.

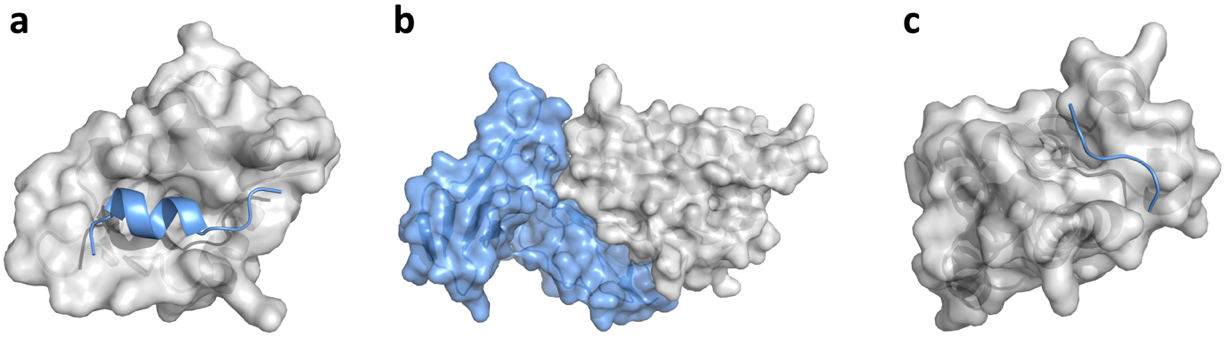

Fig. S8. Three representative PPI target structures. (a) MDM2/p53 exemplifies globular protein-helical peptide interactions. (b) IL-2/IL-2R exemplifies globular protein-globular protein interactions. (c) CREBBP/H3 exemplifies globular protein-peptide interactions.

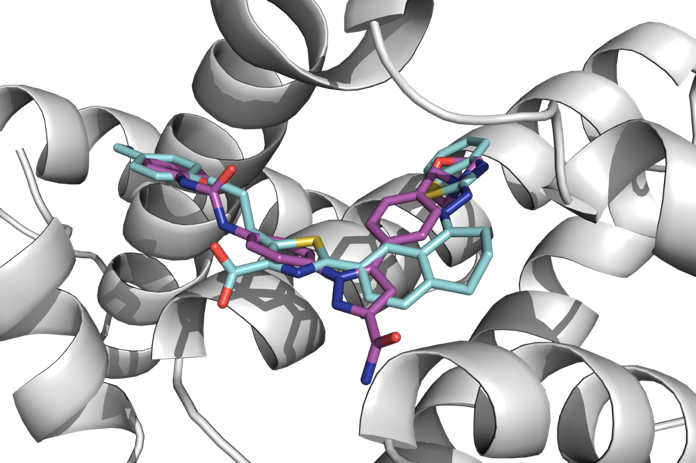

Fig. S9: Overlay of docked poses of the generated ligand (magenta) and WEHI-539 (cyan) bound to BCL-XL (PDB: 3ZLR).

**Supplementary Analyses**

**Design of IL-2/IL-2R inhibitors**

The IL-2/IL-2R interaction plays a critical role in immune regulation, with IL-2 promoting T-cell proliferation through its receptor, IL-2R. Disrupting this interaction offers a potential therapeutic approach for autoimmune diseases and transplant rejection ^1,2^. However, no small-molecule inhibitors targeting IL-2/IL-2R have advanced to clinical trials. In this case, we used a potent inhibitor from the crystal structure (PDB: 1PY2) as the reference ligand ^3^.

Hot2Mol was employed to design inhibitors by mimicking pharmacophores derived from key hot-spot residues on IL-2R—namely Arg36, Leu42, and His120—identified in the crystal structure 1Z92. Then a library of 10,000 molecules was generated and docked against the IL-2 structure (PDB: 1PY2), using the ligand’s position as the docking site. Supplementary Fig. S10 presents the property distribution of the generated molecules in comparison to bioactive compounds targeting the IL-2/IL-2R interaction. The generated molecules exhibit higher QEPPI values, indicating enhanced drug-likeness as PPI inhibitors, and significantly lower SA scores, suggesting better synthetic accessibility. Despite having a lower average molecular weight, these ligands achieved superior docking scores relative to the bioactive molecules (Table S5).

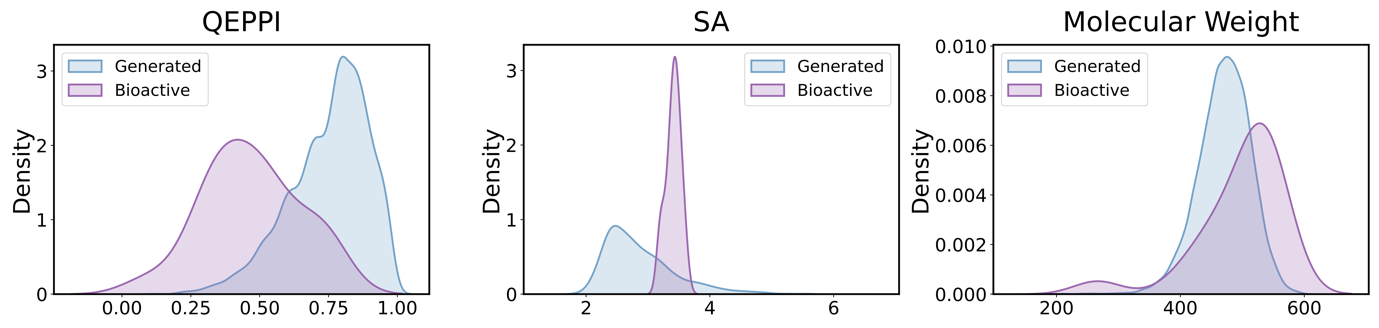

Fig. S10. Property distributions of generated inhibitors and known bioactive molecules targeting IL-2/IL-2R. QEPPI denotes PPI-targeting drug-likeness, SA represents synthetic accessibility, and molecular weight is given in Daltons.

After filtering based on QEPPI scores above 0.5, SA scores below 3.0, and a molecular weight of 600 or less, the molecules were ranked according to their docking scores. From the top ten molecules, two with the best docking poses were selected. PyMOL ^4^ visualization confirmed that these compounds effectively occupy the binding groove on IL-2 (Fig. S11). They achieved docking scores comparable to or better than the reference ligand. Importantly, these higher binding affinities did not compromise synthetic accessibility; the ligands were easier to synthesize and exhibited higher QEPPI scores, indicating their superior drug-likeness. Both generated ligands interacted with key hot-spot residues on IL-2, including Tyr45, Lys43, Phe42, Leu72, Arg38, and Lys35, forming hydrogen bonds, π–π interactions, and salt bridges (Fig. S12).

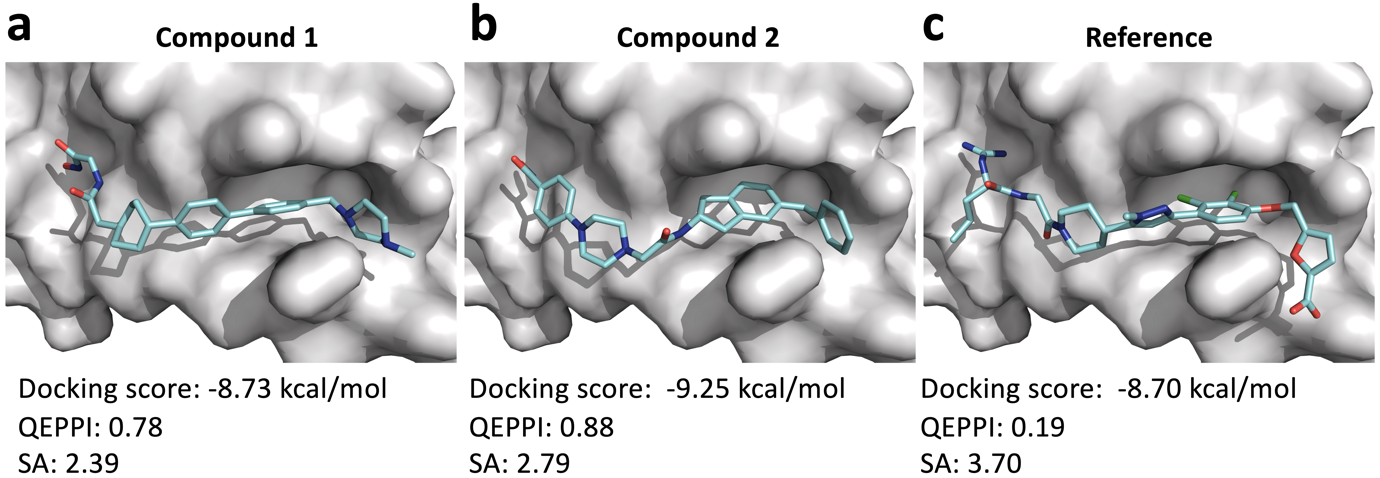

Fig. S11. Docking poses of generated ligands and the reference ligand against the IL-2 structure (PDB: 1PY2). QEPPI represents PPI-targeting drug-likeness, SA is synthetic accessibility, molecular weight is shown in Daltons.

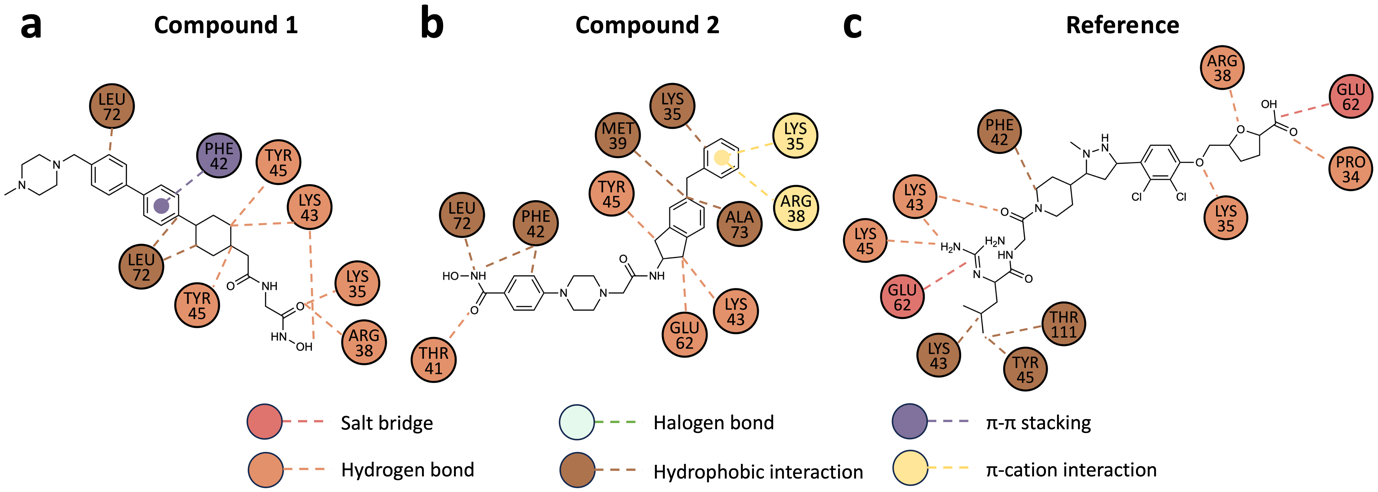

Fig. S12. 2D interaction maps of the generated ligands and the reference ligand against IL-2 structure (PDB: 1PY2). Circles represent amino acid residues, while dashed lines indicate interactions.

**Design of CBP/H3 inhibitors**

The interaction between CBP and histone H3 is vital for chromatin regulation ^5^. Targeting this interaction provides a therapeutic opportunity for treating cancers and other diseases linked to epigenetic dysregulation. Inobrodib (CS1477) is a potent and selective inhibitor for CBP bromodomain inhibitor, which has been used in clinical studies for treating cancers, such as hematologic malignancies ^6^.

Hot2Mol was used to generate 10,000 inhibitors targeting the CBP/H3 interaction by mimicking pharmacophores from key hot-spot residues on the H3 histone—Tyr89, Arg87, and Arg88—identified in the crystal structure 5GH9. These molecules were docked against the CBP structure (PDB:5W0F), with the ligand position from the crystal structure serving as the docking site. Supplementary Fig. S13 presents the property distributions of the generated molecules compared with those of bioactive compounds targeting the CBP/H3 interaction. The generated molecules exhibit higher QEPPI values and lower SA scores, while marginally higher molecular weight.

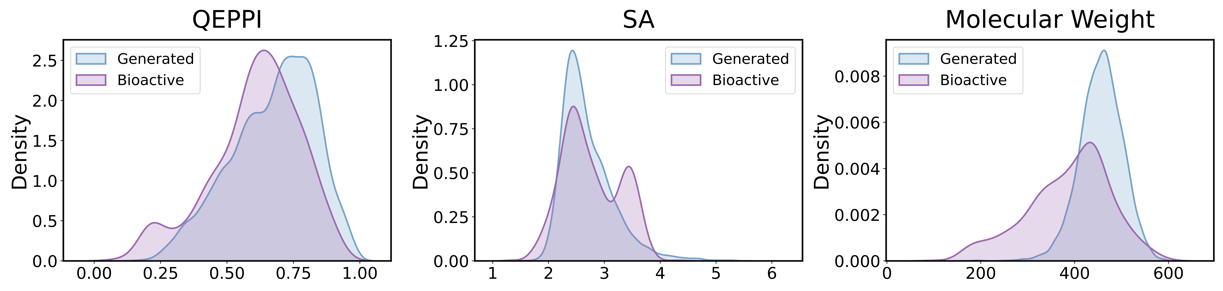

Fig. S13. Property distributions of generated inhibitors and known bioactive molecules targeting CBP/H3. QEPPI denotes PPI-targeting drug-likeness, SA represents synthetic accessibility, and molecular weight is given in Daltons.

The generated molecules were filtered based on QEPPI scores above 0.5, SA scores below 3.0, and molecular weight of 600 or less. These filtered molecules were then ranked according to their docking scores. From the top ten, we selected two with the most favorable docking poses through visual inspection and compared their binding patterns with the Inobrodib. Visualization confirmed that the selected compounds effectively occupy the deep whole in CBP (Fig. S14 a, b), and demonstrate superior docking scores, drug likeness and synthetic accessibility. Both generated ligands interacted with key residues on CBP, including Arg91, Leu38, Val33, Phe29, and Asn86 through hydrophobic interactions or hydrogen bonds (Fig. S14 d-e).

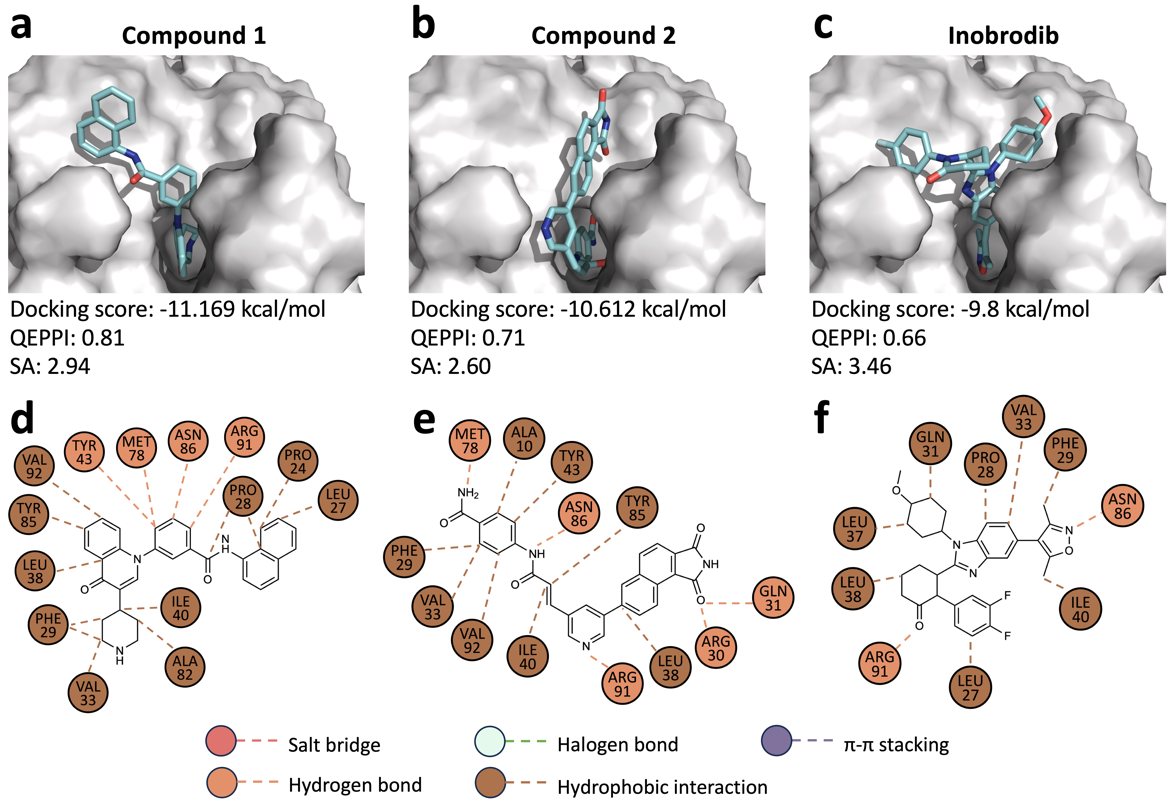

Fig. S14. (a–c) Docking poses of the generated CBP/H3 inhibitors and Inobrodib within the CBP binding site (PDB: 5W0F). QEPPI represents PPI-targeting drug-likeness, SA is synthetic accessibility, molecular weight is shown in Daltons. (d–f) Corresponding 2D interaction maps of the generated ligands and Inobrodib, presented in the same sequence as panels (a–c). Circles represent amino acid residues, while dashed lines indicate interactions.

**Supplementary Tables**

Table S1. Evaluation of molecules generated by Hot2Mol

|  | Validity↑ | Uniqueness↑ | Novelty↑ | Ratio of available molecules↑ | SAscore↓ | QEPPI↑ |
| --- | --- | --- | --- | --- | --- | --- |
| MENIN/MLL | 0.963 | 1.0 | 0.999 | 0.962 | 2.913 | 0.597 |
| IL-2/IL-2R | 0.966 | 0.99 | 1.0 | 0.956 | 2.844 | 0.759 |
| KRAS/SOS1 | 0.982 | 0.928 | 1.0 | 0.911 | 2.537 | 0.58 |
| CREBBP/H3 | 0.977 | 0.992 | 0.998 | 0.967 | 2.675 | 0.672 |
| ZIPA/FTSZ | 0.966 | 0.983 | 0.998 | 0.948 | 2.787 | 0.711 |
| XIAP/CASPASE-9 | 0.984 | 0.936 | 1.0 | 0.921 | 2.563 | 0.564 |
| BCL2/BAX | 0.95 | 0.992 | 0.997 | 0.939 | 2.819 | 0.706 |
| MDM2/P53 | 0.957 | 0.999 | 0.997 | 0.953 | 2.675 | 0.701 |
| TCF4/Beta-catenin | 0.98 | 0.956 | 1.0 | 0.937 | 2.532 | 0.611 |
| BRD4/H4 | 0.985 | 0.95 | 1.0 | 0.936 | 2.576 | 0.64 |

Table S2. Evaluation of molecules generated by PGMG

|  | Validity↑ | Uniqueness↑ | Novelty↑ | Ratio of available molecules↑ | SAscore↓ | QEPPI↑ |
| --- | --- | --- | --- | --- | --- | --- |
| MENIN/MLL | 0.928 | 0.974 | 1.0 | 0.904 | 3.169 | 0.275 |
| IL-2/IL-2R | 0.933 | 0.959 | 1.0 | 0.895 | 3.949 | 0.216 |
| KRAS/SOS1 | 0.867 | 0.997 | 1.0 | 0.864 | 3.74 | 0.543 |
| CREBBP/H3 | 0.981 | 0.965 | 1.0 | 0.947 | 3.762 | 0.464 |
| ZIPA/FTSZ | 0.991 | 0.782 | 0.995 | 0.771 | 2.896 | 0.445 |
| XIAP/CASPASE-9 | 0.969 | 0.996 | 1.0 | 0.965 | 4.262 | 0.567 |
| BCL2/BAX | 0.791 | 0.994 | 0.997 | 0.784 | 3.958 | 0.415 |
| MDM2/P53 | 0.938 | 0.95 | 0.999 | 0.89 | 3.158 | 0.539 |
| TCF4/Beta-catenin | 0.907 | 0.956 | 1.0 | 0.867 | 3.346 | 0.589 |
| BRD4/H4 | 0.905 | 0.964 | 0.997 | 0.869 | 3.238 | 0.492 |

Table S3. Evaluation of molecules generated by Pocket2Mol

|  | Validity↑ | Uniqueness↑ | Novelty↑ | Ratio of available molecules↑ | SAscore↓ | QEPPI↑ |
| --- | --- | --- | --- | --- | --- | --- |
| MENIN/MLL | 1.0 | 0.521 | 1.0 | 0.521 | 2.936 | 0.487 |
| IL-2/IL-2R | 1.0 | 0.453 | 1.0 | 0.453 | 2.714 | 0.574 |
| KRAS/SOS1 | 1.0 | 0.459 | 0.998 | 0.458 | 2.473 | 0.363 |
| CREBBP/H3 | 1.0 | 0.57 | 1.0 | 0.57 | 3.323 | 0.614 |
| ZIPA/FTSZ | 1.0 | 0.518 | 1.0 | 0.518 | 3.301 | 0.517 |
| XIAP/CASPASE-9 | 1.0 | 0.603 | 0.996 | 0.601 | 2.76 | 0.473 |
| BCL2/BAX | 1.0 | 0.632 | 1.0 | 0.632 | 3.145 | 0.571 |
| MDM2/P53 | 1.0 | 0.515 | 1.0 | 0.515 | 3.084 | 0.614 |
| TCF4/Beta-catenin | 1.0 | 0.541 | 1.0 | 0.541 | 3.128 | 0.419 |
| BRD4/H4 | 1.0 | 0.604 | 1.0 | 0.604 | 3.838 | 0.622 |

Table S4. Evaluation of molecules generated by Lingo3DMol

|  | Validity↑ | Uniqueness↑ | Novelty↑ | Ratio of available molecules↑ | SAscore↓ | QEPPI↑ |
| --- | --- | --- | --- | --- | --- | --- |
| MENIN/MLL | 1.0 | 0.995 | 1.0 | 0.995 | 4.076 | 0.276 |
| IL-2/IL-2R | 1.0 | 0.814 | 1.0 | 0.814 | 4.319 | 0.201 |
| KRAS/SOS1 | 1.0 | 0.887 | 1.0 | 0.887 | 3.556 | 0.46 |
| CREBBP/H3 | 1.0 | 0.979 | 0.999 | 0.978 | 3.138 | 0.603 |
| ZIPA/FTSZ | 1.0 | 0.891 | 1.0 | 0.891 | 2.699 | 0.582 |
| XIAP/CASPASE-9 | 1.0 | 0.961 | 1.0 | 0.961 | 3.866 | 0.251 |
| BCL2/BAX | 1.0 | 0.992 | 1.0 | 0.992 | 3.536 | 0.579 |
| MDM2/P53 | 1.0 | 0.961 | 1.0 | 0.961 | 3.831 | 0.623 |
| TCF4/Beta-catenin | 1.0 | 0.887 | 1.0 | 0.887 | 3.306 | 0.409 |
| BRD4/H4 | 1.0 | 0.897 | 1.0 | 0.897 | 2.818 | 0.665 |

Table S5 Docking scores of molecules generated by four DGMs and bioactive compounds

| Target | Hot2Mol | Pocket2Mol | PGMG | Lingo3DMol | Bioactive |
| --- | --- | --- | --- | --- | --- |
| MENIN/MLL | -8.371 ± 0.753 | -6.902 ± 0.785 | -7.078 ± 1.006 | -8.378 ± 0.812 | -7.443 ± 0.927 |
| IL-2/IL-2R | -7.996 ± 0.763 | -7.110 ± 1.161 | -5.970 ± 1.058 | -6.165 ± 1.164 | -7.292 ± 0.867 |
| KRAS/SOS1 | -8.540 ± 0.796 | -6.497 ± 0.754 | -8.295 ± 0.743 | -7.061 ± 0.781 | -6.929 ± 0.835 |
| CREBBP/H3 | -8.936 ± 0.776 | -8.527 ± 1.040 | -6.995 ± 0.912 | -7.888 ± 0.865 | -7.962 ± 1.033 |
| ZIPA/FTSZ | -7.554 ± 0.674 | -7.064 ± 0.969 | -6.918 ± 0.679 | -6.670 ± 0.829 | -7.304 ± 0.489 |
| XIAP/CASPASE-9 | -7.443 ± 0.821 | -6.133 ± 0.851 | -7.387 ± 0.644 | -6.623 ± 0.670 | -6.622 ± 0.606 |
| BCL2/BAX | -9.187 ± 0.869 | -8.751 ± 1.344 | -7.841 ± 0.918 | -8.552 ± 1.048 | -8.599 ± 0.975 |
| MDM2/P53 | -8.343 ± 0.726 | -8.428 ± 1.045 | -8.286 ± 0.718 | -8.179 ± 0.784 | -8.150 ± 0.707 |
| TCF4/Beta-catenin | -6.736 ± 0.669 | -5.879 ± 0.683 | -6.798 ± 0.717 | -5.903 ± 0.843 | -6.874 ± 0.627 |
| BRD4/H4 | -8.766 ± 0.727 | -8.888 ± 1.041 | -7.815 ± 0.999 | -7.867 ± 0.855 | -7.998 ± 0.873 |

Table S6. Results of the ablation study

|  | Target | Validity↑ | Uniqueness↑ | Novelty↑ | Ratio of available molecules↑ | SAscore↓ | QEPPI↑ | Mean Docking Scores↑ |
| --- | --- | --- | --- | --- | --- | --- | --- | --- |
| original | MENIN/MLL | 0.963 | 1.0 | 0.999 | 0.962 | 2.913 | 0.597 | -8.326 |
|  | IL-2/IL-2R | 0.966 | 0.99 | 1.0 | 0.956 | 2.844 | 0.759 | -7.996 |
|  | KRAS/SOS1 | 0.982 | 0.928 | 1.0 | 0.911 | 2.537 | 0.58 | -8.38 |
|  | CREBBP/H3 | 0.977 | 0.992 | 0.998 | 0.967 | 2.675 | 0.672 | -8.936 |
|  | ZIPA/FTSZ | 0.966 | 0.983 | 0.998 | 0.948 | 2.787 | 0.711 | -7.554 |
|  | XIAP/CASPASE-9 | 0.984 | 0.936 | 1.0 | 0.921 | 2.563 | 0.564 | -7.449 |
|  | BCL2/BAX | 0.95 | 0.992 | 0.997 | 0.939 | 2.819 | 0.706 | -9.188 |
|  | MDM2/P53 | 0.957 | 0.999 | 0.997 | 0.953 | 2.675 | 0.701 | -8.343 |
|  | TCF4/Beta-catenin | 0.98 | 0.956 | 1.0 | 0.937 | 2.532 | 0.611 | -6.701 |
|  | BRD4/H4 | 0.985 | 0.95 | 1.0 | 0.936 | 2.576 | 0.64 | -8.768 |
| without-EGNN | MENIN/MLL | 0.992 | 0.768 | 0.999 | 0.761 | 2.899 | 0.374 | -8.183 |
|  | IL-2/IL-2R | 0.986 | 0.672 | 1.0 | 0.663 | 3.32 | 0.671 | -7.212 |
|  | KRAS/SOS1 | 0.998 | 0.456 | 1.0 | 0.455 | 2.475 | 0.497 | -7.682 |
|  | CREBBP/H3 | 0.982 | 0.742 | 1.0 | 0.729 | 2.962 | 0.588 | -8.018 |
|  | ZIPA/FTSZ | 0.999 | 0.692 | 0.999 | 0.69 | 2.902 | 0.579 | -7.613 |
|  | XIAP/CASPASE-9 | 0.997 | 0.417 | 1.0 | 0.416 | 2.485 | 0.514 | -6.806 |
|  | BCL2/BAX | 0.99 | 0.814 | 1.0 | 0.806 | 3.909 | 0.515 | -8.679 |
|  | MDM2/P53 | 0.99 | 0.794 | 1.0 | 0.786 | 3.024 | 0.643 | -8.235 |
|  | TCF4/Beta-catenin | 0.998 | 0.491 | 1.0 | 0.49 | 2.432 | 0.514 | -6.553 |
|  | BRD4/H4 | 0.998 | 0.552 | 1.0 | 0.551 | 2.684 | 0.456 | -8.089 |
| without-property-constraint | MENIN/MLL | 0.976 | 0.995 | 1.0 | 0.971 | 2.851 | 0.428 | -9.092 |
|  | IL-2/IL-2R | 0.976 | 0.996 | 0.999 | 0.971 | 3.063 | 0.596 | -8.020 |
|  | KRAS/SOS1 | 0.841 | 0.992 | 1.0 | 0.834 | 2.816 | 0.398 | -8.466 |
|  | CREBBP/H3 | 0.96 | 0.999 | 1.0 | 0.959 | 3.224 | 0.582 | -8.764 |
|  | ZIPA/FTSZ | 0.985 | 0.995 | 1.0 | 0.98 | 2.759 | 0.619 | -7.684 |
|  | XIAP/CASPASE-9 | 0.818 | 0.994 | 1.0 | 0.813 | 3.03 | 0.354 | -7.475 |
|  | BCL2/BAX | 0.975 | 0.998 | 1.0 | 0.973 | 2.769 | 0.637 | -9.473 |
|  | MDM2/P53 | 0.977 | 0.998 | 1.0 | 0.975 | 2.821 | 0.586 | -8.732 |
|  | TCF4/Beta-catenin | 0.944 | 0.995 | 1.0 | 0.939 | 2.732 | 0.448 | -6.896 |
|  | BRD4/H4 | 0.904 | 0.99 | 1.0 | 0.895 | 2.959 | 0.407 | -8.756 |
